# *In vivo* dissection of Rhoa function in vascular development using zebrafish

**DOI:** 10.1101/2021.03.27.437282

**Authors:** Laura M. Pillay, Joseph J. Yano, Andrew E. Davis, Matthew G. Butler, Keith A. Barnes, Vanessa L. Reyes, Daniel Castranova, Aniket V. Gore, Matthew R. Swift, James R. Iben, Amber N. Stratman, Brant M. Weinstein

## Abstract

**Rationale:** The small monomeric GTPase RHOA acts as a master regulator of signal transduction cascades by activating effectors of cellular signaling, including the Rho-associated protein kinases ROCK1/2. Previous *in vitro* cell culture studies suggest that RHOA can regulate many critical aspects of vascular endothelial cell (EC) biology, including focal adhesion, stress fiber formation, and angiogenesis. However, the specific *in vivo* roles of RHOA during vascular development and homeostasis are still not well understood.

**Objective:** In this study we examine the *in vivo* functions of RHOA in regulating vascular development and integrity in zebrafish.

**Methods and Results:** We use zebrafish RHOA-ortholog (*rhoaa*) mutants, transgenic embryos expressing wild type, dominant-negative, or constitutively active forms of *rhoaa* in ECs, and a pharmacologic inhibitor of ROCK1/2 to study the *in vivo* consequences of RHOA gain- and loss-of-function in the vascular endothelium. Our findings document roles for RHOA in vascular integrity, developmental angiogenesis, and vascular morphogenesis.

**Conclusions:** Our results indicate that either too much or too little RHOA activity leads to vascular dysfunction *in vivo*.

## INTRODUCTION

An endothelial cell (EC) monolayer forms the innermost layer of all blood vessels. This layer acts as a physical barrier to maintain blood vessel integrity and selective permeability, regulating the exchange of oxygen, solutes, nutrients, and immune cells between the blood and tissues. Under normal physiologic conditions, the EC barrier is maintained throughout the development of both embryonic and adult vascular networks, which form predominantly by angiogenesis – the “sprouting” of new vessels from pre-existing ones. Within the brain, disruption of the EC barrier leads to inflammation, edema, or hemorrhagic stroke, producing a spectrum of disease that includes neurologic dysfunction, disability, cognitive impairment, or death ^1^. The formation, maintenance, and disassembly of EC barrier junctions are therefore highly regulated processes under strict spatiotemporal control ^2^.

There are two principal types of barrier-forming EC junctions, tight junctions and adherens junctions ^2, 3^. Tight junctions are continuous inter-EC junctions that limit paracellular permeability by controlling the exchange of cells, fluids, and solutes between the vessel lumen and surrounding tissues. Adherens junctions serve as anchors that maintain stable cellular associations under mechanical stress, or dynamic adhesive connections that facilitate cellular movement (e.g., during angiogenic blood vessel growth or remodeling). Both junction types are tethered to the actin cytoskeleton, and their assembly, maintenance and remodeling are modulated by actin-myosin contractile forces ^4^. Although the intracellular signaling pathways that control EC cytoskeletal remodeling and junctional adhesion are still an active area of investigation, previous studies have implicated the Ras Homology (Rho)-family member RHOA in these processes (reviewed in ^2, 4–7^).

RHOA is a small, monomeric GTPase that transduces stimuli from hormones, growth factors, cytokines, and transmembrane proteins to downstream effectors of cellular signaling. RHOA acts as a molecular switch, cycling between a GDP-bound inactive state and a GTP-bound active state ^8, 9^. In response to upstream cellular signals, guanine nucleotide exchange factors (GEFs) switch RHOA to its active, GTP-bound state ^10^. Active RHOA can then activate effectors including the Rho-associated protein kinases ROCK1 and ROCK2 ^11^. GTPase-activating proteins (GAPs) inactivate RHOA by restoring it to its inactive, GDP-bound state ^10^. Although previous studies suggest RHOA may regulate many critical aspects of vascular EC biology including cellular migration, proliferation, actin stress fiber formation, focal adhesion, and adherens junction dynamics ^7, 12–16^, the effects of RHOA on cytoskeletal dynamics and intercellular junction formation appear to be context-specific. Junction-destabilizing factors such as VEGF or thrombin ^17–20^, and junction-stabilizing factors such as Angiopoietin 1 or Sphingosine-1-phosphate ^21–24^ both appear to require RHOA and ROCK.

Due to the early embryonic lethality of *Rhoa*-knockout mice ^25^ most of the functional characterization of RHOA in the vascular endothelium to date has been performed in cell culture, by overexpressing dominant negative or constitutively active forms of RHOA or by treating ECs *in vitro* with exogenous factors that modulate their growth and development ^4^. ECs in culture are also typically grown under static growth conditions in the absence of other cell types. ECs *in vivo* are subject to continuous or pulsatile flow, producing laminar shear stress and cyclic stretch — forces that have been shown to differentially impact RHOA subcellular localization, signaling, and its effects on cytoskeletal remodeling ^26, 27^. ECs *in vivo* also interact with numerous other cell types that influence their behavior, including vascular smooth muscle cells, pericytes, macrophages, astrocytes, and neurons ^28^. Consequently, although *in vitro* cell culture studies have been useful for identifying the cellular activities of RHOA in ECs, the *in vivo* biological significance of these studies remains unclear.

Studies of human patients and disease models with cerebral cavernous malformations (CCMs) have provided evidence that RHOA may play a role in regulating vascular integrity *in vivo*. CCMs are clusters of abnormally dilated blood vessels that exhibit vascular hyperpermeability and are extremely prone to rupture. Familial cases of CCM are associated with mutations in CCM complex genes (CCM1, 2, or 3) ^29^. The ECs of murine CCM models exhibit elevated RHOA and ROCK1/2 activity, along with a concomitant increase in actin stress fiber formation and reduced junctional integrity ^30–32^. Treatment of murine CCM models with ROCK1/2 inhibitor decreases their hemorrhage and lesion burden ^31, 33–35^. CCM studies therefore suggest that RHOA signaling may negatively regulate vascular integrity *in vivo*. In support of this hypothesis, another study demonstrated that loss of EC RHOA in postnatal Cdh5-Cre^ERT2^*; Rhoa^f/f^* mice is sufficient to block vascular leakage following exogenous histamine but not VEGF-treatment ^36^.

Other *in vitro* studies have also suggested roles for RHOA and ROCK1/2 in angiogenesis. RHOA and ROCK1/2 are required for VEGF-stimulated capillary formation from ECs in both *in vitro* cell culture and in mouse *ex vivo* retinal explants ^15, 37, 38^. However, studies examining *in vivo* requirements for RHOA in developmental angiogenesis have produced conflicting reports. In one study, EC-specific deletion of *Rhoa* in Tie2-Cre; *Rhoa^f/f^* mice generated an embryonic lethal phenotype with vascular hypoplasia, yolk sac vascular remodeling defects, and dilated blood vessels, suggesting that EC-specific RHOA is required for proper vascular development and embryonic survival ^39^. In contrast, a more recent study by a different group failed to observe any gross vascular abnormalities in their Tie2-Cre; *Rhoa^f/f^* EC-specific RHOA deficiency mouse model ^40^. However, viable Tie2-Cre; *Rhoa^f/f^* pups were obtained at sub-Mendelian ratios in this study. It is therefore possible that EC RHOA deficiency produced an incompletely-penetrant, early embryonic-lethal phenotype in this model. Overall, existing studies of EC-specific RHOA function have provided limited, contradictory information on the requirement of EC RHOA in regulating embryonic vascular development *in vivo*.

Zebrafish provide a powerful model organism for studying embryonic vascular development and homeostasis. Forward and reverse genetic approaches in zebrafish have identified molecular pathways and genes that regulate vascular integrity and development and have clear orthologues in mammals ^41^. Embryos develop externally, making them highly amenable to genetic manipulation. Optical transparency and transgenic strains permit visualization of hemorrhage and vascular development *in vivo*, in real-time through time-lapse microscopy ^42, 43^. The small size of zebrafish embryos allows them to survive and continue their development for many days in the absence of circulatory function, facilitating assessment of the specificity of vascular defects.

In this study, we use zebrafish as a model organism to assess the *in vivo* functions of Rhoa and its effectors Rock1/2 in regulating vascular development and integrity. We examine zebrafish with mutations in the zebrafish RHOA-ortholog *rhoaa*, transgenic animals with EC-specific expression of wild type or mutant forms of *rhoaa*, and animals treated with a pharmacological inhibitor of Rock1/2, using high-resolution optical imaging to study the consequences of Rhoa gain- and loss-of function in the vascular endothelium. Our results suggest that cranial vessel integrity in developing zebrafish is highly sensitive to either increased or decreased Rhoa gene dosage and activity, and that Rhoa and Rock1/2 are required for proper embryonic blood vessel growth and patterning *in vivo*.

## METHODS

### Data availability

All data and materials reported in this manuscript are available from the Zebrafish International Resource Center (https://zebrafish.org) or upon request by contacting the corresponding author.

### Animal care and pharmacological treatments

Fish were housed in a large zebrafish dedicated recirculating aquaculture facility (4 separate 22,000L systems) in 6L and 1.8L tanks. Fry were fed rotifers and adults were fed Gemma Micro 300 (Skretting) once per day. Water quality parameters were routinely measured, and appropriate measures were taken to maintain water quality stability (water quality data available upon request). Zebrafish husbandry and research protocols were reviewed and approved by the NICHD Animal Care and Use Committee, in accordance with the Guide for the Care and Use of Laboratory Animals of the National Institutes of Health. The facility is accredited by the Association for Assessment and Accreditation of Laboratory Animal Care (AAALAC). EK strain zebrafish were used for all experiments unless otherwise noted. Embryos were grown at 28.5°C in embryo media (EM) or methylene blue-containing fish water (2 mL of 0.1% methylene blue per 1L media) and staged according to standardized morphological criteria ^44^. EM was supplemented with 0.006% 1-phenyl 2-thiourea (PTU) (Sigma), to prevent pigment formation in post-24 hours post fertilization (hpf) embryos. Rockout (Rho Kinase Inhibitor III; Santa Cruz Biotechnology) was used to inhibit Rock1/2 activity ^45^. Rockout was dissolved in Dimethyl sulfoxide (DMSO) to generate a stock concentration of 30 mM and then diluted to a working concentration in EM + PTU. Embryos were treated from 24 hpf onward with 25 μM or 50 μM Rockout or equivalent dilutions of DMSO (solvent controls) in EM + PTU. All embryos were raised at 28.5°C and imaged or assessed for phenotypes at 52 hpf.

### Whole mount *in situ* hybridization

Examination of gene expression by whole mount *in situ* hybridization was performed essentially as previously described ^46, 47^, with the following modifications: Riboprobes were designed to hybridize primarily to the 3’UTR of each gene to minimize cross-reactivity between zebrafish orthologues with coding sequence similarities (see **Table S1** for primer sequences used to amplify riboprobes). Prior to mRNA *in situ* hybridization analyses, embryos were fixed in 4% paraformaldehyde (PFA)/phosphate-buffered saline (PBS) overnight at 4°C or 4–5 hours at room temperature (RT) with gentle agitation on a rotating platform. Embryos were permeabilized in 10 μg/ml proteinase K for 5 minutes (24–32 hpf embryos), or 30 minutes (48 – 52 hpf embryos) at RT.

### Transgenic zebrafish lines

Published transgenic fish lines used in experiments include *Tg*(*gata1*:*DsRed*)^sd2Tg^ ^48^, *Tg(kdrl:EGFP)^s8^*^43 49^, *Tg(fli:EGFP)^y1^* ^50^, and *Tg(egfl7:GAL4FF)^gSAlzGFFD478A^* ^51, 52^.

All cloning was performed using SLiCE cloning techniques unless otherwise noted ^53^. See **Table S1** for primer sequences used for SLiCE cloning. The p14xUAS-Tom-2A plasmid was made by cloning the Tomato fluorescent protein (Tom) and P2A viral cleavage peptide (2A) ^54, 55^ sequences into the multiple cloning site (MCS) of pT1ump-14xUAS-MCS-POUT (pT1UMP), which includes an E1b basal promoter, the Ocean pout antifreeze protein polyA, and is flanked by Tol1 arms ^56^. Select *rhoaa* coding sequences were PCR-amplified from wild type or *rhoaa*-mutant (R17, Δ3) zebrafish cDNAs and cloned into pCRII-TOPO (Invitrogen), according to manufacturer’s specifications. To make p14XUAS-Tom-2A-rhoaa, p14XUAS-Tom-2A-rhoaaR17, and p14XUAS-Tom-2A-rhoaaΔ3, *rhoaa* coding sequences were PCR-amplified from wild type or *rhoaa*-mutant / pCRII-TOPO template and cloned into p14xUAS-Tom-2A. p14XUAS-Tom-2A-rhoaaN19 and p14XUAS-Tom-2A-rhoaaV14 were made by site-directed mutagenesis of p14XUAS-Tom-2A-rhoaa. G0 *Tg(UAS-Tom-2A)*, *Tg(UAS-Tom-2A-rhoaa)*, *Tg(UAS-Tom-2A-rhoaaR17)*, *Tg(UAS-Tom-2A-rhoaaΔ3)*, *Tg(UAS-Tom-2A-rhoaaN19)*, and *Tg(UAS-Tom-2A-rhoaaV14)* transgenic zebrafish embryos were each generated by injecting individual p14xUAS-Tom-2A or p14xUAS-Tom-2A-rhoaa isoforms into zebrafish embryos along with zebrafish-optimized Tol1 transposase RNA ^57^.

### Mutant zebrafish lines and genotyping analyses

The *y172* (aka *R17*, *rhoaa^R17^,* or *rhoaa^y172^*) mutant was identified in an F2 ENU mutagenesis screen for dominant mutants displaying a hemorrhage phenotype at 2-4 dpf in the *Tg(fli:EGFP)^y1^* background (**Figure S1**). Bulked segregant analysis and fine genetic mapping of *y172* were carried out as described previously ^58^ and linked *y172* to a 3.148 MB region on Chromosome 8, in between determined SSLP markers ZC221E5-6 and ZK83F4-7 (**Figure S1**). Whole genome sequencing (Illumina HiSeq)-based identification of deleterious mutations within the critical mapping region revealed a p.G17R mutation in *rhoaa*. The *rhoaa^y172^* or *rhoaa^R17^* mutants were genotyped using derived cleaved amplified polymorphic sequence (dCAPS) analysis ^59–61^. Briefly, genomic DNA was isolated from adults or embryos as previously described ^62^. PCR was used to introduce an XbaI restriction enzyme site in R17-mutant, but not WT *rhoaa* PCR product, using rhoaaY172Geno-F and rhoaaY172Geno-R primers (**Table S1**). PCR products were cleaved with XbaI, generating 194 bp WT and/or 169 bp R17-mutant products, analyzed by gel electrophoresis.

*rhoaa^Δ3^* mutants were generated using CRISPR/Cas9 technology as previously described ^63^, using the single-guide RNA sequence ATCGTTGGAGATGGAGCCTGTGG, which was selected using CHOPCHOP (https://chopchop.cbu.uib.no) ^64^ to target the Rhoaa GTP/GDP binding domain. *rhoaa^Δ3^* mutant embryos and adults were genotyped as previously described ^65^. Genomic DNA was extracted using the REDExtract-N-Amp Tissue PCR Kit (Sigma): 50 µl of extraction solution and 12.5 µl of tissue preparation solution was added to each embryo. Samples were mixed by vortex, incubated 10 min at RT, vortexed, and heated to 95°C for 4 min. 50 µl of neutralization solution B was added, samples were vortexed, and stored at −80°C or used immediately. Solution volumes were doubled for DNA extraction from adult tail clips. Gene-specific, tailed genotyping primers (rhoaaM13Ftail-F and rhoaaPigtail-R, **Table S1**) were designed to amplify a 184 bp fragment spanning the CRISPR binding site. A 10 µl total volume AmpliTaq-Gold (Life Technologies) PCR reaction was set up for each sample, containing 1 µl 10X PCR Gold Buffer, 0.5 µl 25 mM MgCl2, 1 µl 0.5 mM rhoaaM13Ftail-F primer, 1 µl 1 mM rhoaaPigtail-R primer, 0.2 µl 10 mM FAM-M13 primer, 0.1 µl 10 mM each dNTP Master Mix, 0.1 µl TaqGold polymerase, 1 µl of a 1:10 dilution of genomic DNA in water, and 5.1 µl water. PCR conditions were as follows: 10 min denaturation at 95°C; 35 cycles of 95°C for 30 sec, 58°C for 30 sec, and 72°C for 30 sec; and 10 min final extension at 72°C. 2 µl of PCR product was combined with 0.2 µl ROX400HD (Life Technologies) and 9.8 µl HiDi Formamide, and the solution was denatured at 95°C for 5 min and run on a ABI3130XL Genetic Analyzer (Applied Biosystems), using GeneMapper software (Life Technologies).

### Endothelial cell culture

Passage 1-6 human umbilical vein endothelial cells (HUVEC, Lonza) were cultured in 0.003% Endothelial cell growth supplement (Millipore), 0.01% Heparin, 20% FBS, 2.5 ug/mL Amphotericin B and 0.1X penicillin/streptomycin in M199 media (Gibco) on 1 mg/ml gelatin coated tissue culture flasks at 37 °C in a humidified 5% CO2 incubator, as previously described ^66^.

Lentiviral constructs for use in HUVEC *in vitro* were generated using SLiCE cloning techniques unless otherwise noted ^53^. See **Table S1** for primer sequences. Constructs with the CMV promoter were used because the PGK promoter has previously been shown to exhibit reduced or absent activity in endothelial cells, including HUVEC ^67, 68^. The sCMV promoter was amplified from pCS2+ ^69^ by PCR and cloned into the pCDH-EF1α-MCS-(PGK-GFP-T2A-PURO) bidirectional promoter lentiviral vector (System Biosciences), replacing the PGK promoter, to generate pCDH-EF1α-MCS-(sCMV-GFP-T2A-PURO). Wild type or mutant (*R17, N19, V14, Δ3*) *rhoaa* coding sequences were PCR-amplified from p14XUAS-Tom-rhoaa constructs and cloned into the MCS of pCDH-EF1α-MCS-(sCMV-GFP-T2A-PURO) to generate pCDH-rhoaa (WT, R17, N19, V14, Δ3). Tomato (Tom) fluorescent protein sequences were then PCR-amplified and cloned into the MCS of pCDH-EF1α-MCS-(sCMV-GFP-T2A-PURO) or into pCDH-rhoaa to generate pCDH-Tom control or pCDH-Tom-rhoaa constructs. Notably, previous studies indicate that N-terminal epitope or fluorophore-tagging does not significantly impact RHOA subcellular localization or activity in endothelial cells ^70, 71^.

Lentivirus was made from pCDH-Tom and pCDH-Tom-rhoaa (WT, R17, Δ3, N19, or V14) using the ViraPower Lentiviral Expression System (Invitrogen): 9 µg ViraPower Packaging Mix and 3 µg plasmid were added to 1.5 mL Opti-MEM. 36 µl Lipofectamine 2000 was added to 1.5 mL Opti-MEM. Solutions were gently mixed, incubated 5 minutes (min) at room temperature (RT), then combined, gently mixed, and incubated 20 min at RT. The resulting solution was added dropwise to a T75 flask of 90% confluent HEK293T cells grown in 10% FBS in DMEM media (Gibco) and gently mixed by tilting. Cells were incubated at 37 °C in a humidified 5% CO2 incubator, and inspected for syncytia formation and Tomato/GFP-fluorescence. Cell media was replaced daily. Lentivirus-containing media was collected at both 2 and 3 days post-transfection: Cell debris was pelleted by centrifugation 15 min at 3,000 rpm and discarded. Virus-containing media was filtered through a Millex-HA 0.45 µm filter and stored at −80 °C until use.

Stable HUVEC lines expressing Tomato, Tomato-rhoaa, Tomato-rhoaa*^R17^*, Tomato-rhoaa^Δ3^, or Tomato-rhoaa^N19^ from integrated lentiviral vectors were generated by infecting HUVEC with the appropriate Lentivirus: 2.5 µl 6 mg/mL Polybrene was mixed with 500 µl of HUVEC growth media. Media was removed from 90% confluent HUVEC grown in 6-well tissue culture plates, and Polybrene mixture was added dropwise to cells. 1 mL of Lentivirus-containing media was added to culture, and HUVEC were grown at 37 °C until greater than 30% expressed GFP. Transgene-expressing HUVEC were selected by Puromycin treatment. We were unable to recover stable lines of HUVEC expressing Tomato-rhoaa^V14^ due to non-viability of cells.

### Immunohistochemistry

HUVEC were transferred to 1.33% collagen / phosphate-buffered saline (PBS)-coated Lab-Tek chamber slides (ThermoFisher) at 90% confluency and allowed to attach overnight at 37°C. Cells were fixed for 5 min in 100% methanol at RT. All washes and incubations were performed with gentle agitation on a rotating platform at RT, unless otherwise indicated. Cells were washed 4 x 5 min in tris-buffered saline (TBS; 136 mM NaCl, 25 mM Tris, pH 7.4), then 15 min in tris-glycine buffer (0.025M Tris, 0.192M glycine, pH 8.3). Cells were permeabilized 30 min in 0.1% Saponin/TBS, washed 4 x 5 min in TBS, and blocked 1 hour in TBS + 0.1% Tween-20 (TBST) + 2.5 % bovine serum albumin (BSA) + 5% sheep serum, then incubated in at 1:500 dilution of anti-VE Cadherin antibody (abcam, ab33168) in blocking solution, overnight at 4 °C. Cells were washed 4 x 5 min in TBST, incubated 1 hour in a 1:1000 dilution of Alexa Fluor 488-conjugated secondary antibody (ThermoFisher) in blocking solution, then incubated in a 1:2000 dilution of Hoechst 33342/TBS solution (Invitrogen) for 20 min. Cells were washed 4 x 5 min in TBST, mounted in Fluorescent Mounting Medium (Dako), and stored at 4°C in the dark.

### Western analysis

Zebrafish crude protein lysates were prepared by adding 1 mL cold deyolking buffer (110 mM NaCL, 5.5 mM KCl, 10 mM Tris pH 8.5, 2.7 mM CaCl2) containing cOmplete Mini Protease Inhibitor Cocktail (Roche) to 20 – 30 embryos. Yolk was disrupted by gentle pipetting followed by 30 sec of 1100 rpm vortex. Tissue was pelleted by centrifuging 1 min at 3,000 rpm and supernatant was discarded. 0.7 mL of cold wash buffer (110 mM NaCl, 5.5 mM KCl, 10 mM Tris pH 8.5, 2.7 mM CaCl2) containing protease inhibitor was added. Tissues were subjected to Dounce homogenization, centrifuged 5 min at 5,000 rpm and the supernatant discarded. 5 µl of 2X LDS Sample Buffer containing 5 % 2-Mercaptoethanol per embryo was added to each sample. Lysates were vortexed, incubated 10 min at 85°C, then stored at −20°C.

Immediately prior to analysis, a 1:2 dilution of lysate to 1X LDS Sample Buffer was prepared and heated 10 min at 85°C. 15 µl of diluted lysate was run out on NuPAGE Novex Bis-Tris gels using the XCell Sure Lock Mini-Cell system (Invitrogen) at 140 V, according to manufacturer’s specifications. Proteins were transferred to methanol-treated PVDF membranes using a Mini Trans-Blot Cell wet-transfer apparatus (BIO-RAD), according to manufacturer’s specifications at 400 mA for 1.5 hours. All membrane washes and incubations were performed with gentle rocking. Following transfer, membranes were washed 5 min in Tris-buffered saline with 0.1% Tween-20 (TBST), then allowed dry completely. Membranes were rehydrated in methanol, rewashed with TBST, and blocked at least 1 hr in 5% skim milk in TBST. Membranes were incubated in primary antibody diluted in TBST (1:500 anti-RHOA antibody, Cytoskeleton ARH04) or diluted in block (1:30,000 anti-a-alpha tubulin antibody, Sigma T6199) overnight at 4 °C. After 5 x 5 min RT TBST washes, membranes were incubated in 1:2000 digital HRP-conjugated secondary antibody (Kindle Bioscience) in TBST. After TBST washes, a 1-Shot Digital ECL reaction (Kindle Bioscience) was performed, and images were acquired using a KwikQuant Imager (Kindle Bioscience). Quantification of relative band density was performed using ImageJ software ^72^. Data is reported as the percent average density from a minimum of 3 blots from independent experiments ± standard deviation.

### Microscopy and imaging

Fluorescent images of HUVEC were acquired using a Zeiss LSM 880 confocal microscope with Fast Airyscan, using a 63X objective with optical zoom. All images were acquired using the same settings.

Whole-mount *in situ* hybridization images and transmitted-light images of live zebrafish embryos were collected using a Leica M205 stereo microscope and LAS V4.7 software, with extended depth of focus z-stack image processing. Fluorescent images of zebrafish embryos were collected using a Nikon Yokogawa CSU-W1 spinning disk confocal microscope. Transmitted light movies of live zebrafish embryos were collected using a Leica M205 stereo microscope and LAS V4.7 software, or a Nikon Yokogawa CSU-W1 spinning disk confocal microscope. Prior to imaging, live embryos were immobilized in EM containing buffered MS-222 and embedded in 0.8% low melting point agarose in EM. Zebrafish embryos were genotyped after imaging and data analysis.

All fluorescent images were processed using ImageJ ^72^ or Imaris (Oxford Instruments) and all figures were assembled using Powerpoint (Microsoft) and/or Photoshop (Adobe). Copies of original transmitted light images of live zebrafish embryos were globally adjusted for red color saturation to enhance red blood cell visibility equally across matched (control and treatment) samples. Movies were assembled using Imaris (Oxford Instruments), Premiere Pro (Adobe), and After Effects (Adobe).

### Statistical analyses

For each zebrafish experiment, data from two or more independent biological replicates with separate cohorts of embryos were analyzed for the main effects of treatment.

For analyses of hemorrhage in *rhoaa^R17^* mutant embryos, homogeneity across replicates was determined using the *G*-test of independence, and homogenous datasets (heterogeneity *G*-value ≥ 0.05) were combined for statistical analysis. Heterogeneous replicate datasets were analyzed separately and combined only when statistical analyses yielded identical results for each replicate dataset. Significant differences among treatments were determined using two-tailed Fisher’s exact tests on cumulative raw counts, with Bonferroni correction applied to multiple comparisons (alpha = 0.05).

For analyses of wild type versus *rhoaa^Δ3/+^* incross progeny, embryos with a specific phenotype were categorized (wild type, hemorrhaging, no circulation; wild type CtAs, reduced CtAs, overbranching CtAs) and counted. Homogeneity across replicates was determined using the *G*-test of independence, and homogenous datasets (heterogeneity *G*-value ≥ 0.05) were combined for statistical analysis. Heterogeneous replicate datasets were analyzed separately and combined only when statistical analyses yielded identical results for each replicate dataset. Significant differences among treatments were determined using Fisher-Freeman-Halton exact tests of independence on raw counts (alpha = 0.05).

For EC-specific transgenic Tomato or *rhoaa* isoform overexpression experiments and Rockout treatment experiments, significant phenotypic differences among groups or treatments across biological replicates were determined using the Cochran-Mantel-Haenszel test for repeated 2×2 tests of independence (alpha = 0.05).

## RESULTS

### Zebrafish *rhoaa* mutants with distinct cranial vascular integrity and patterning defects

We identified a novel mutant *(y172)* in a forward-genetic ENU mutagenesis screen for dominant or partially-dominant zebrafish mutants with intracranial hemorrhage (**Figure S1A,B**). Genetic fine mapping and genome sequencing suggested that the causative defect in the *y172* mutant is a Glycine to Arginine substitution at amino acid 17 in the evolutionarily conserved GTP/GDP-binding domain^73, 74^ of *rhoaa,* a zebrafish orthologue of mammalian RHOA (**Figure 1A**; **Figure S1C-F**). We therefore subsequently refer to *y172* mutants as *rhoaa^R17^* or R17 mutants. Whole-mount *in situ* hybridization analyses show that *rhoaa* is broadly expressed in wild type embryos at 26 hours post fertilization (hpf), with particularly strong expression in the head (**Figure 1B**), becoming almost exclusively expressed in the head by 48 hpf (**Figure 1C,D**). Starting at approximately 48 hpf, both R17/+ heterozygous and R17/R17 homozygous mutant embryos begin to exhibit incompletely penetrant cranial hemorrhage (**Figure 1E-G,M**) with variable expressivity ranging from small hematomas barely visible under a light microscope to large hemorrhages that fill one or more brain ventricles (**Figure S1B**). To determine if R17 mutants have altered blood vessel morphology or patterning, we crossed them to a *Tg(kdrl:EGFP)^s843^, Tg*(*gata1*:*DsRed*)*^sd2Tg^* double transgenic reporter line with EGFP-labeled endothelial cells (ECs) and DsRed-labeled primitive blood cells ^48, 49^. By 52 hpf, both R17/+ and R17/R17 embryos develop vasculogenic primary cranial blood vessels, including the lateral dorsal aortae (LDA) and primordial hindbrain channels (PHBC), as well as angiogenic cranial central arteries (CtA), although CtA growth is sometimes slightly delayed in R17/R17 animals (**Figure 1I-L**)^75^. R17/+ and R17/R17 embryos also exhibit normal development of trunk blood vessels, including the dorsal aorta (DA), cardinal vein (CV), and intersegmental vessels (ISVs) at 52 hpf (**Figure S2A-C**).

**Figure 1.**
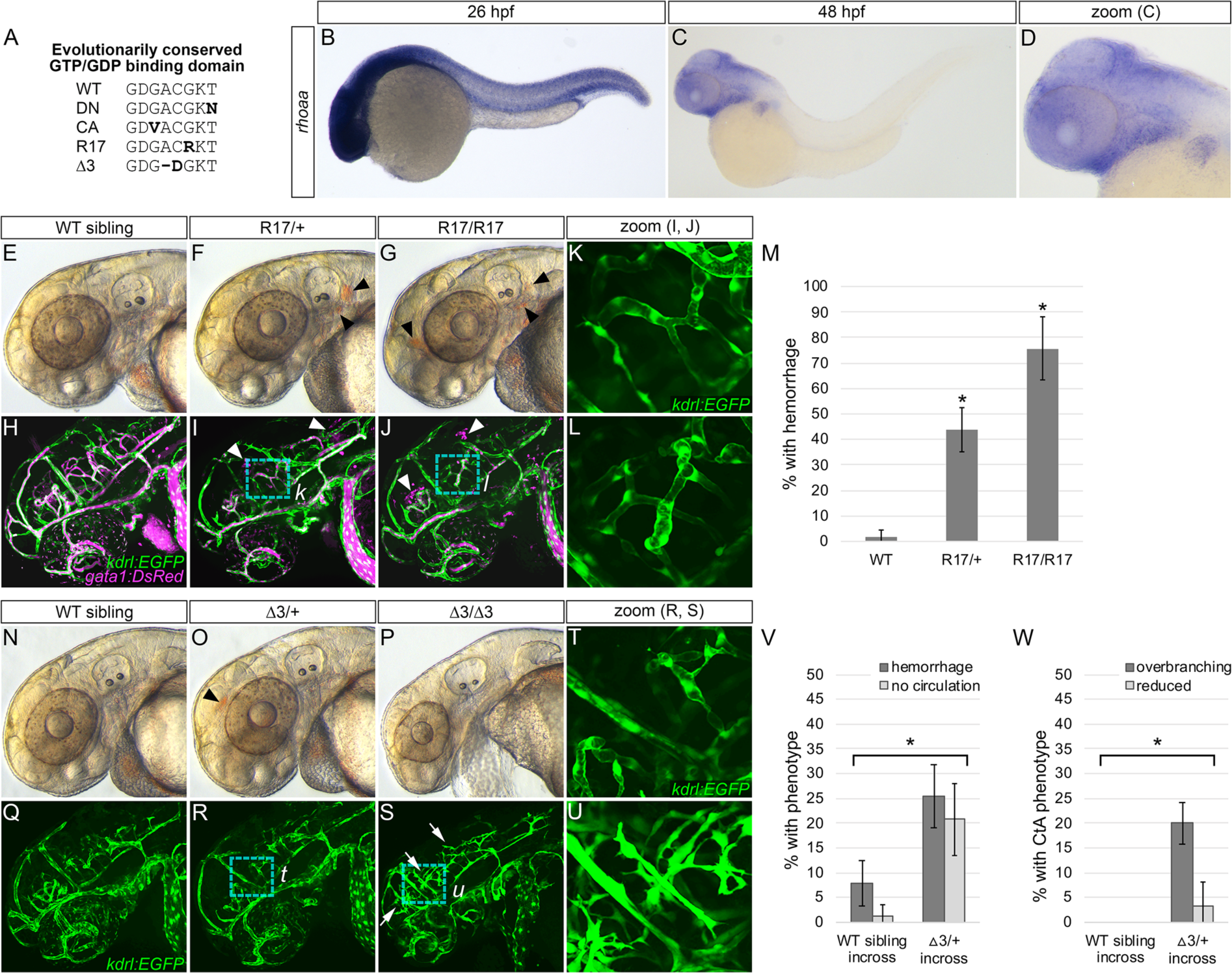
*rhoaa* mutant zebrafish embryos exhibit vascular integrity and patterning defects. (**A**) Partial amino acid sequences of wild type (WT) Rhoaa, known dominant negative (DN) Rhoaa^N19^, known constitutively active (CA) Rhoaa^V14^, Rhoaa^R17^, Rhoaa^△^*^3^*, and Rhoaa^△^*^6^* GTP/GDP-binding domains. (**B**) Whole-mount *in situ* hybridization analysis of *rhoaa* gene expression in a 26 hours post fertilization (hpf) wild type (WT) zebrafish embryo shown in lateral view, anterior to left. (**C**) Whole-mount *in situ* hybridization analysis of *rhoaa* gene expression in a 48 hpf WT zebrafish embryo shown in lateral view, anterior to left. (**D**) Magnified view of head in C. (**E-G**) Stereoscope transmitted light images of 52 hpf *Tg(kdrl:EGFP; gata1:DsRed*); *rhoaa^R17/+^* incross progeny heads shown in lateral view, anterior to left. Arrowheads indicate hemorrhage. (**H-J**) Confocal images of the vasculature (*kdrl:EGFP*; green) and primitive erythrocytes (*gata1:DsRed*; magenta) in the same animals as in E-G. Arrowheads indicate hemorrhage. (**K,L**) Zoomed-in view of central arteries (CtAs; boxed areas in J and K). (**M**) Quantitation of the percentage of embryos of indicated genotype with hemorrhage by 52 hpf. \**P* < 0.0003 compared to WT siblings, by Fisher’s exact test with Bonferroni correction. (**N-P**) Stereoscope transmitted light images of 52 hpf *Tg(kdrl:EGFP*); *rhoaa^△3/+^* incross progeny. Arrowhead indicates hemorrhage. (**Q-S**) Confocal images of the vasculature (*kdrl:EGFP*; green) of the same animals as in O-Q. Arrows indicate overbranching CtAs. (**T,U**) Zoomed-in view of CtAs (boxed area in R and S). (**V,W**) Quantitation of the percentage of WT sibling incross progeny (left two columns) versus *rhoaa^△3/+^* incross progeny (right two columns) that either (V) have hemorrhage or lack circulation at 52 hpf (\**P* ?? 0.0024 compared to WT incross progeny, by Fisher-Freeman-Halton test), or (W) have reduced sprouting or overbranching of CtAs (\**P* < 0.001, compared to WT incross progeny, by Fisher-Freeman-Halton test). Mean ± SD is shown for each graph.

The *rhoaa* gene is one of five highly-conserved zebrafish RHOA orthologues (**Figure S3A,B**) exhibiting highly overlapping developmental expression patterns (**Figure S3C-H**). The strong possibility of compensation by other zebrafish *rhoa* orthologues and the dominant nature of the R17 mutation suggest that R17 may be acting as either a dominant negative (DN) or constitutively active (CA) allele of *rhoaa.* In support of this hypothesis, previous studies have shown that mutating the RHOA GTP/GDP-binding domain yields both DN and CA RHOA alleles. For example, RHOA^V14^ is a CA version of human RHOA that is unable to hydrolyze GTP, while RHOA^N19^ is a DN version of human RHOA that preferentially remains bound to GDP ^73, 76^ (**Figure 1A**). To help further explore the effects of and validate the R17 mutation, we used CRISPR/Cas9 mutagenesis ^63^ to generate an additional zebrafish mutant with a small, in-frame 3-base pair deletion and amino acid substitution in the Rhoaa GTP/GDP-binding domain, (designated *rhoaa^Δ3^* or Δ3; **Figure 1A**). We found that the R17 and Δ3 mutants exhibit partially overlapping but distinct phenotypes (**Figure 1, Figure S2**).

Like R17/+ heterozygotes, Δ3/+ heterozygous embryos exhibit wild type morphology with incompletely penetrant cranial hemorrhage beginning at 48 hpf (**Figure 1O,V**) and normal cranial and trunk blood vessel patterning at 52 hpf (**Figure 1R, Figure S2E**). However, unlike R17/R17 animals, most Δ3/Δ3 homozygotes lack blood circulation and exhibit heart edema, despite having a heartbeat (**Movie S1; Figure 1P,V**). They also have smaller heads with atypical morphology (**Figure 1P**). Although the vasculogenic primary cranial vessels (LDA, PHBC) are normally patterned in Δ3/Δ3 homozygous mutants, they exhibit ectopic growth and branching of the angiogenic central arteries (CtA) and, as might perhaps be expected from the lack of circulation, their vessels are poorly dilated (**Figure 1S,U,W**). Overall trunk blood vessel patterning is normal in 52 hpf Δ3/Δ3 homozygotes, although these vessels (especially the ISVs) also fail to fully dilate (**Figure S2F**). Together, these data suggest that while R17/+ and Δ3/+ heterozygotes both exhibit impaired cranial vascular integrity, Δ3/Δ3 homozygotes have additional vascular defects not found in R17/R17 homozygotes.

The difference between R17/R17 and Δ3/Δ3 homozygotes does not appear to be the result of differences in the relative amounts of Rhoa proteins, since both show very similar (approximately 50%) reductions in total Rhoa, as measured by western blotting of whole-embryo zebrafish protein lysates collected at 50 hpf (**Figure S4**). These results suggest that the Rhoaa^R17^ and Rhoaa^Δ3^ proteins may instead possess distinct activities.

### Rhoaa alleles have differential effects on adherens junctions in cultured primary ECs *in vitro*

Vascular permeability is modulated primarily by the integrity of EC-EC adherens and tight junction complexes^2^. Adherens junctions consist of transmembrane cadherins, such as Cdh5/VE-Cadherin. Previous studies using primary human umbilical vein ECs (HUVEC) *in vitro* have shown that increased RHOA activity disrupts adherens junctions (Haidari et al., 2011; Wojciak-Stothard et al., 2002), while dominant-negative suppression of RHOA signaling does not alter the distribution of adherens junction proteins in unstimulated cells (Wójciak-Stothard et al., 2001). To examine the effects of expressing different mutant *rhoaa* proteins on EC-EC junctions *in vitro,* we used lentiviral transduction to generate stable HUVEC lines expressing either wild type *rhoaa*, known dominant-negative *rhoaa^N19^*, *rhoaa^R17^*, or *rhoaa^Δ3^*. We also attempted to generate stable HUVEC lines expressing known constitutively-active *rhoaa^V14^*, but cells transduced with this allele are not viable (data not shown). We examined adherens junction morphology in confluent monolayers of stable *rhoaa* isoform-expressing HUVEC lines using immunohistochemical staining with an anti-Cdh5 antibody (**Figure 2**).

**Figure 2.**
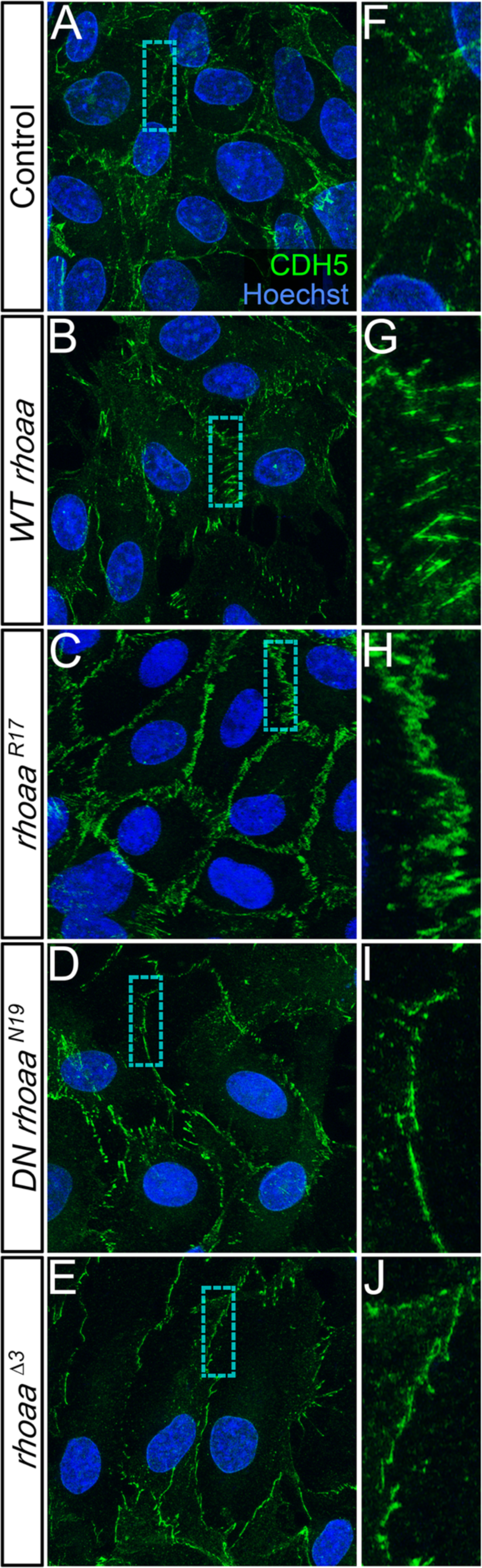
Rhoaa isoforms exhibit distinct activities *in vitro.* **(A-E)** Left: anti-CDH5 immunostaining of adherens junctions (green) in HUVEC stably transfected with wild type (WT) or mutant zebrafish *rhoaa* isoforms. Nuclei labeled with Hoescht 33342 (blue). **(F-J)** Zoomed-in view of boxed areas shown in panels on left.

Compared to “empty-vector” transduced controls (**Figure 2A,F**), HUVEC expressing WT *rhoaa* exhibit jagged, highly discontinuous junctional Cdh5 staining (**Figure 2B,G**), as previously reported for ECs with excess RHOA pathway activation (Pronk et al., 2017). HUVEC expressing *rhoaa^R17^* also exhibit jagged junctional staining (**Figure 2C,H**), although in this case, the staining is much less discontinuous than in HUVEC expressing WT *rhoaa*. In contrast, HUVEC expressing either the known dominant-negative *rhoaa^N19^* (**Figure 2D,I**) or *rhoaa^Δ3^* (**Figure 2E,J**) exhibit thin, mostly continuous junctional Cdh5 staining.

Together these results suggest that the Rhoaa^R17^ protein may have reduced or weak dominant-negative activity, while Rhoaa^Δ3^ likely acts as a strong dominant-negative allele of Rhoaa.

### Vascular defects are associated with either increased and decreased Rhoaa levels or activity in the zebrafish vascular endothelium *in vivo*

To further explore the *in vivo* consequences of increased or decreased Rhoa activity specifically in the vascular endothelium, we generated a series of *Tg(UAS:Tomato-2A-rhoaa)* transgenic zebrafish co-expressing Tomato fluorescent protein together with either wild type *rhoaa*, the *rhoaa^R17^* or *rhoaa^Δ3^* mutant isoforms, or known dominant-negative or constitutively-active *rhoaa* isoforms (*DN rhoaa^N19^* or *CA rhoaa^V14^*) under Gal4 control. We crossed these germline *UAS:Tomato-2A-rhoaa* transgenics to *Tg(egfl7:GAL4ff); Tg(UAS:EGFP)* double-transgenic zebrafish with vascular EC-specific Gal4 and EGFP reporter expression (**Figure 3A**).

**Figure 3.**
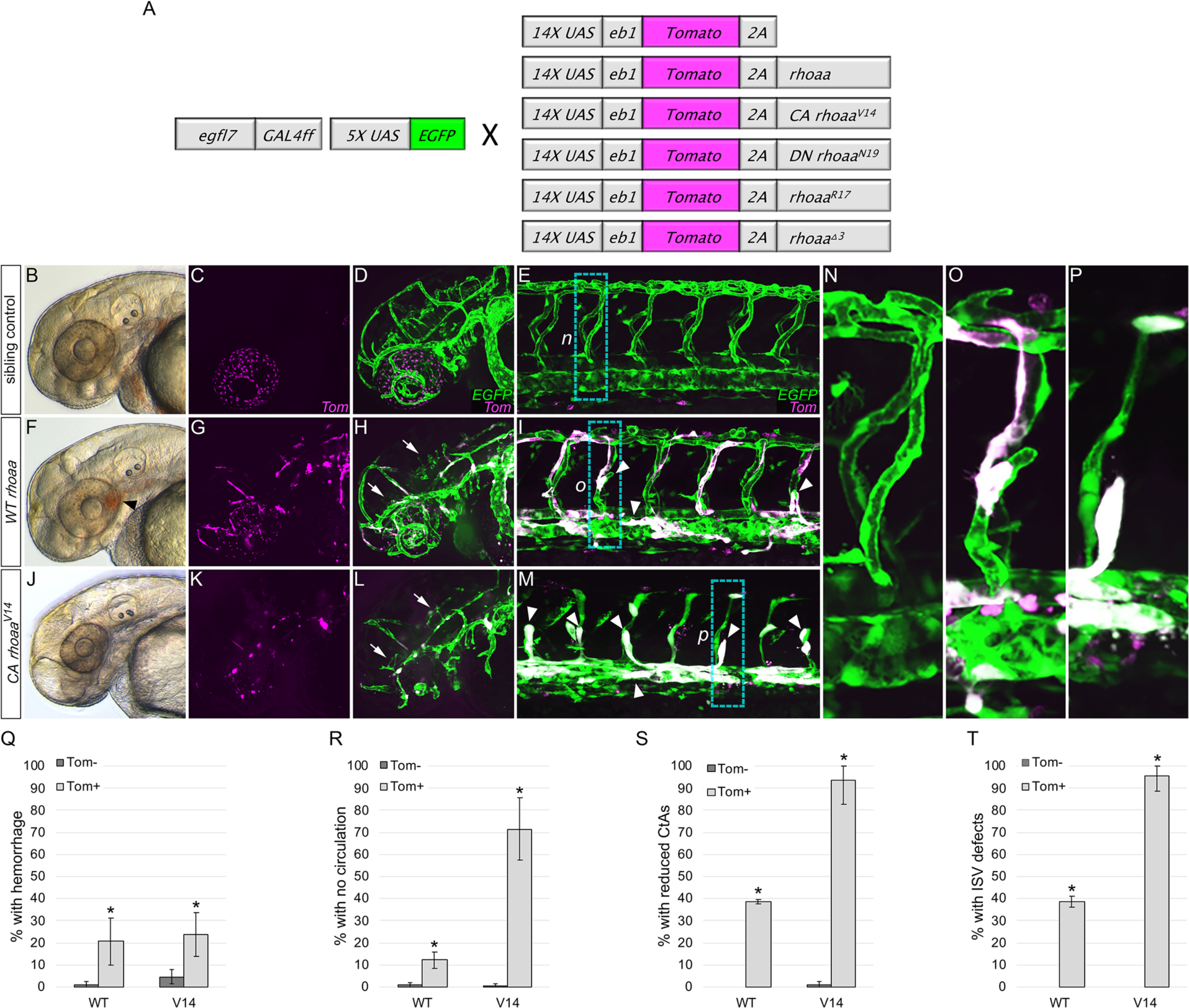
Increased endothelial cell Rhoa activity induces vascular integrity and patterning defects *in vivo.* (**A**) Schematics of transgenics used for conditional wild type or mutant *rhoaa* expression in zebrafish endothelial cells. (**B,F,J**) Stereoscope transmitted light images of 52 hpf progeny derived from a cross between *Tg(egfl7:GAL4ff), Tg(UAS:EGFP)* and *Tg(UAS:Tomato-2A-rhoaa)* (B,F) or *Tg(UAS:Tomato-2A-rhoaa^V14^)* (J) fish. Panel B shows a non-Tomato/*rhoaa* transgene-expressing control sibling of the Tomato/WT *rhoaa* transgene-expressing animal in panel F. Black arrowhead in F indicates hemorrhage. (**C,D,G,H,K,L**) Tomato (C,G,K) and Tomato/EGFP (D,H,L) confocal images of cranial endothelial cells in the same embryos as in panels B,F,J. Arrows indicate reduced CtA sprouting. (**E,I,M**) Tomato/EGFP confocal images of trunk endothelial cells in 52 hpf progeny derived from a cross between *Tg(egfl7:GAL4ff), Tg(UAS:EGFP)* and *Tg(UAS:Tomato-2A-rhoaa)* (E,I) or *Tg(UAS:Tomato-2A-rhoaa^V14^)* (M) fish. White arrowheads indicate ISV sprouting defects or impaired vessel dilation. (**N-P**) Magnified Tomato/EGFP confocal images of trunk vasculature in the boxed areas in panels E,I,M. (**V-Y**) Quantitation of the percentage of 52 hpf Tomato-positive WT Rhoaa-expressing or Rhoaa^V14^-expressing (light grey columns) animals or their Tomato-negative siblings (dark grey columns) with specified phenotype. Phenotypes shown in the graphs are cranial hemorrhage (Q), lack of circulation (R), reduced cranial CtA sprouting (S), or defective trunk ISV sprouting (T). \**P* ?< 0.0001 compared to Tomato-negative sibling controls, by Cochran-Mantel-Haenszel test. Mean ± SD is shown for each graph.

EC-specific expression of Tomato alone has no effect on vascular integrity or development in zebrafish embryos (**Figure S5**). Compared to their control Tomato-negative siblings (**Figure 3B-E,N,Q-T**), embryos expressing ectopic *WT rhoaa* in ECs exhibit incompletely penetrant cranial hemorrhage (**Figure 3F,Q**), lack of circulation (**Figure 3R**), reduced growth of cranial CtAs (**Figure 3G,H,S**), impaired growth of trunk ISVs (**Figure 3I,O,T**), and vessel dilation defects (**Figure 3G-I,O**). Embryos expressing *CA rhoaa^V14^* in ECs also sometimes exhibit cranial hemorrhage (**Figure 3Q**) and have much more penetrant and severe circulation, CtA/ISV angiogenesis, and blood vessel dilation defects (**Figure 3K-M,P,R-T, Movie S2**) than either WT-*rhoaa* expressing animals or their control Tomato-negative siblings (**Figure 3R-T, Figure S6K-N,S**). Unlike WT-*rhoaa* expressing ECs, *CA rhoaa^V14^*-expressing ECs also frequently have a rounded, punctate morphology and/or appear fragmented, particularly in *rhoaa^V14^*-expressing cranial ECs (**Figure 3K-M, Movie S3**). Combined, these results suggest that increased Rhoaa dosage and/or activity in zebrafish ECs produces both vascular integrity and vascular growth defects, with high levels of Rhoa activation (in *CA rhoaa^V14^*-expressing ECs) inducing rounded, punctate EC morphology and/or EC fragmentation.

To study the consequences of decreased Rhoaa activity the vascular endothelium *in vivo*, we examined transgenic zebrafish embryos with EC-specific expression of known dominant-negative *DN rhoaa^N19^.* Compared to Tomato-negative sibling controls (**Figure 4A-D,Q,U,V**) embryos expressing *DN rhoaa^N19^* in ECs exhibit severe cranial hemorrhage (**Figure 4E,U**) and reduced CtA vessel growth at 52 hpf (**Figure 4F,G,V**). Unlike cranial CtAs expressing EC-specific *DN rhoaa^N19^*, most trunk ISVs expressing this isoform grow well and successfully reach the DLAV by 52 hpf, although a subset of ISVs are abnormally dilated and/or detach from the DA, CV, or DLAV (**Figure 4H,R, Movie S4**). Like embryos expressing *DN rhoaa^N19^* in the vascular endothelium, compared to Tomato-negative sibling controls (**Figure 4U,V**, **Figure S7K-N,S**), embryos with EC-specific *rhoaa^Δ3^* expression exhibit severe cranial hemorrhage, reduced CtA vessel growth, and abnormally dilated and/or detached ISVs (**Figure 4I-L,S,U,V**), supporting the idea that Rhoaa^Δ3^ is acting as a dominant-negative Rhoaa protein.

**Figure 4.**
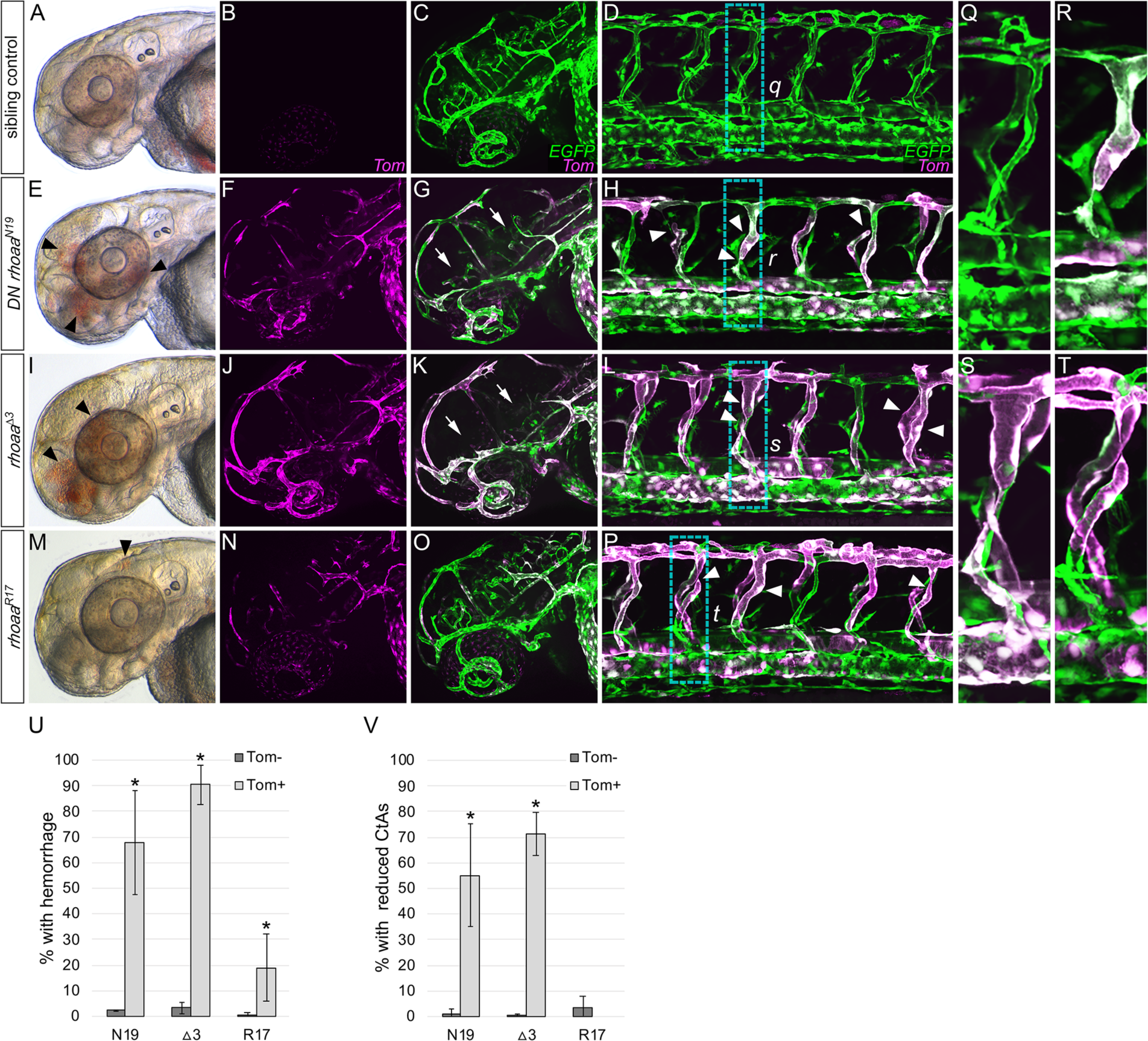
Endothelial cell specific expression of dominant negative *rhoaa* induces cranial vessel integrity, patterning, and trunk vessel dilation defects *in vivo*. (**A,E,I,M**) Stereoscope transmitted light images of 52 hpf progeny derived from a cross between *Tg(egfl7:GAL4ff), Tg(UAS:EGFP)* and *Tg(UAS:Tomato-2A-rhoaa^N19^)* (A,E)*, Tg(UAS:Tomato-2A-rhoaa^△3^)* (I), or *Tg(UAS:Tomato-2A-rhoaa^R17^*) (M) fish. Panel A shows a non-Tomato/*rhoaa* transgene-expressing control sibling of the Tomato/*rhoaa^N19^* transgene-expressing embryo in panel E. Black arrowheads in E indicate hemorrhage. (**B,C,F,G,J,K,N,O**) Tomato (B,F,J,N) and Tomato/EGFP (C,G,K,O) confocal images of cranial endothelial cells in the same embryos as in panels A,E,I,M. Arrows indicate reduced CtA sprouting. (**D,H,L,P**) Tomato/EGFP confocal images of trunk endothelial cells in 52 hpf progeny derived from a cross between *Tg(egfl7:GAL4ff), Tg(UAS:EGFP)* and *Tg(UAS:Tomato-2A-rhoaa^N19^)* (D,H)*, Tg(UAS:Tomato-2A-rhoaa^△3^)* (L), or *Tg(UAS:Tomato-2A-rhoaa^R17^*) (P) fish. Arrowheads indicate ISV dilation, detachment, or growth defects. (**Q,R,S,T**) Magnified Tomato/EGFP confocal images of trunk vasculature in the boxed areas in panels D,H,L,P. (**U,V**) Quantitation of the percentage of Tomato-positive embryos (light grey columns) or their Tomato-negative siblings (dark grey columns) with cranial hemorrhage (U) or reduced cranial CtA sprouting (V) at 52 hpf. \**P* ?< 0.0001 compared to Tomato-negative sibling controls, by Cochran-Mantel-Haenszel test. Mean ± SD is shown for each graph.

In contrast, compared to Tomato-negative sibling controls (**Figure 4U,V, Figure S7U-X,C’**) zebrafish embryos with transgene-driven EC-specific expression of the Rhoaa^R17^ protein show relatively low levels of cranial hemorrhage and no reduction in CtA growth (**Figure 4M-O,U,V**). A small subset of these embryos do exhibit abnormally dilated and/or detached trunk ISVs (**Figure 4P,T**), although this phenotype is not as severe or penetrant as seen in *DN rhoaa^N19^* (**Figure 4H,R**) or *DN rhoaa^Δ3^* (**Figure 4L,S**) expressing animals. These results suggest that Rhoaa^R17^ may be a very weak, dominant negative isoform of Rhoaa compared to DN Rhoaa^N19^ or Rhoaa^Δ3^.

### Rock function is required for cranial angiogenesis and to maintain vascular integrity *in vivo*

The Rho-associated protein kinases ROCK1 and ROCK2 are some of the best characterized RHOA effectors^11^. Whole-mount *in situ* hybridization analyses of the zebrafish ROCK orthologues *rock1* and *rock2a* revealed that their expression is enriched in the head at 52 hpf, in nearly identical patterns to *rhoaa* (**Figure 5A-D**), making them attractive candidates to mediate RHOA signaling in regulating cranial blood vessel growth and integrity. To determine whether loss of Rock1/2 activity results in comparable vascular defects to *rhoaa^Δ3/Δ3^* mutants or animals with EC-specific expression of *rhoaa^Δ3^*, we treated zebrafish from 24 - 52 hpf with Rockout, a cell-permeable pharmacological inhibitor of Rock1/2 kinase activity previously used in zebrafish ^45, 77, 78^. This treatment interval was selected to bypass earlier gastrulation defects caused by Rock inhibition ^79^. Rockout-treated animals exhibit dose-dependent, highly penetrant cranial hemorrhage and CtA growth defects at 52 hpf (**Figure 5E-N**).

**Figure 5.**
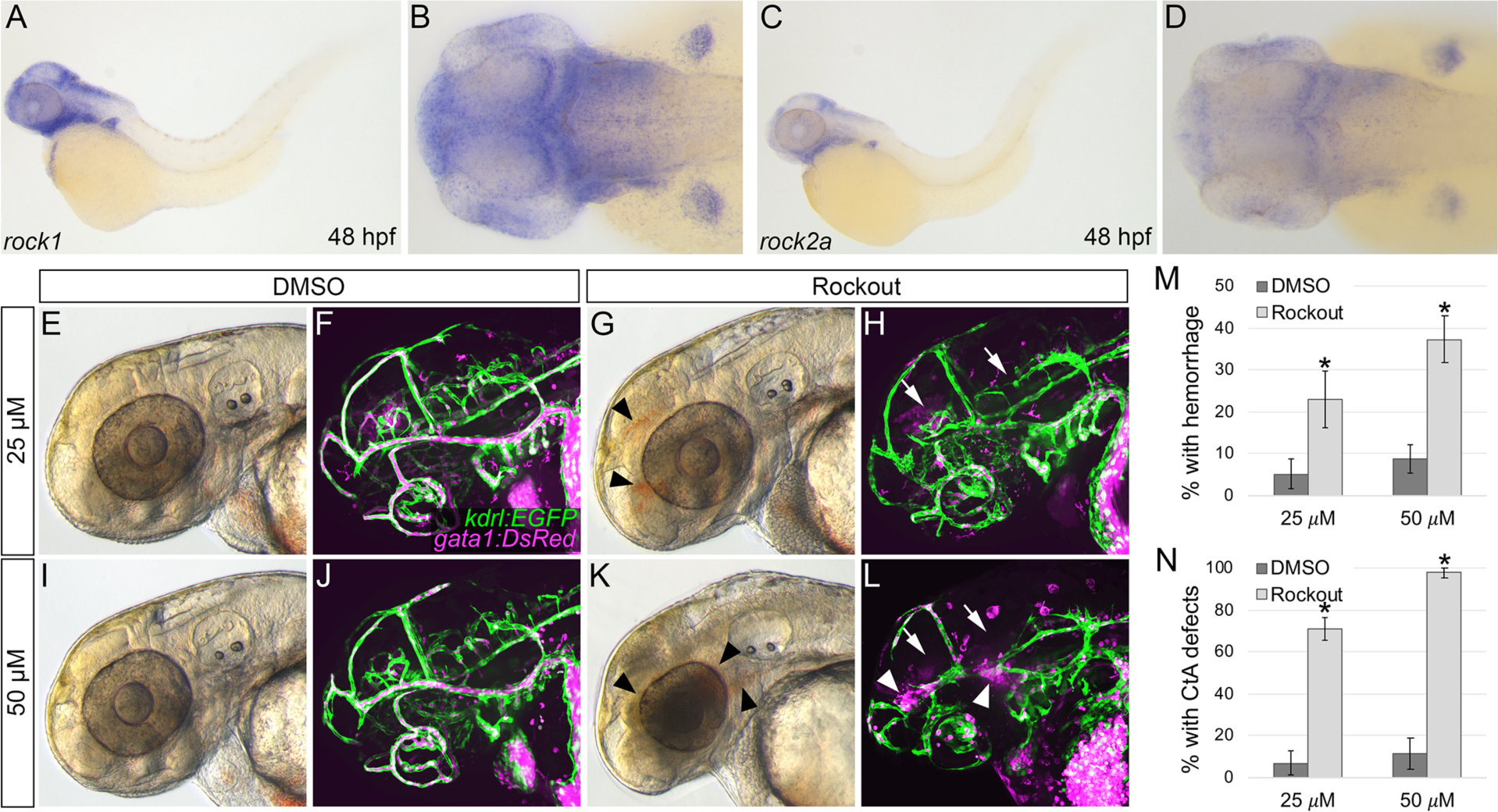
Pharmacological inhibition of Rhoa effectors Rock1/2 produces cranial hemorrhage and cranial angiogenesis defects. (**A-D**) Whole-mount *in situ* hybridization analyses of 48 hpf wild type zebrafish embryos probed for *rock1* (A,B) or *rock2a* (C,D) gene expression. (A,C) Lateral views, anterior to left. (B,D) Higher magnification dorsal views of the heads shown in panels A,C. (**E,G,I,K**) Stereoscope transmitted light images of DMSO-treated control sibling (E,I) or 25 μM (G) or 50 μM (K) Rock1/2 inhibitor (Rockout)-treated 50 hpf *Tg(kdrl:EGFP), Tg(gata1a:DsRed)* embryos. Arrowheads indicate hemorrhage. (**F,H,J,L**) Confocal images of cranial vessels (EGFP-positive, green) and blood cells (DsRed-positive, magenta) in the same embryos shown in panels E,G,I,K. Arrows indicate CtA defects, arrowheads indicate hemorrhage. (**M,N**) Quantitation of the percentage of 52 hpf Rockout-treated (light grey columns) or DMSO-treated (dark grey columns) sibling embryos with cranial hemorrhage (M) or cranial CtA defects (N). \**P* < 0.0001 compared to DMSO-treated controls, by Cochran-Mantel-Haenszel test. Mean ± SD is shown for each graph.

## DISCUSSION

In this study, we have examined the function of the small, monomeric GTPase Rhoa in regulating vascular development and integrity *in vivo.* We used zebrafish RHOA-ortholog (*rhoaa*) mutants, transgenic embryos expressing wild type, dominant-negative, or constitutively active forms of *rhoaa* in ECs, and a pharmacologic inhibitor of the Rhoa effectors Rock1/2 to study the *in vivo* developmental consequences of Rhoa gain- and loss-of-function in the vascular endothelium. Our findings document roles for Rhoa function in vascular integrity, developmental angiogenesis, and vascular morphogenesis *in vivo*.

### Identification and characterization of novel zebrafish mutant alleles of Rhoa

We isolated and characterized two mutants carrying novel alleles of the zebrafish RHOA-orthologue *rhoaa (rhoaa^?^*^3^ and *rhoaa^R17^;* **Figure 1**). Both mutants display partially dominant cranial hemorrhage phenotypes as heterozygotes. However, while *rhoaa^?^*^3/^*^?^*^3^ homozygotes fail to develop circulation and exhibit both cranial vascular patterning and morphological defects, *rhoaa^R17/R17^* homozygotes exhibit wild type circulation, vascular patterning, and morphology. Both mutants have comparable levels of total Rhoa protein, suggesting that their phenotypic differences reflect differences in Rhoaa isoform activities, not protein levels. This idea is supported by the differential phenotypes of ECs expressing *rhoaa^R17^* or *rhoaa^?3^* both *in vitro* in primary human ECs (**Figure 2**) and *in vivo* in zebrafish embryos (**Figure 4**).

Human umbilical vein endothelial cells (HUVEC) expressing either the known dominant-negative (DN) *rhoaa^N19^* or *rhoaa^?3^* show essentially identical adherens junction morphology, while HUVEC expressing *rhoaa^R17^* show an intermediate phenotype between cells expressing wild type *rhoaa* and cells expressing *DN rhoaa^N19^* or *rhoaa^?3^* (**Figure 2**). Similarly, zebrafish with transgene-driven EC-specific expression of either *DN rhoaa^N19^* or *rhoaa^?3^* exhibit severe cranial hemorrhage, a strong reduction in angiogenic cranial vessel growth, and dilation/detachment of trunk vessels. *rhoaa^R17^*-expressing zebrafish embryos display far less penetrant cranial hemorrhage and trunk vessel dilation/detachment phenotypes, and no reduction in cranial vessel growth (**Figure 4**). In contrast to the results with these other mutants, the punctate EC morphologies and EC fragmentation observed in the known constitutive-active *rhoaa^V14^* mutant suggest that this mutant may be incompatible with endothelial cell viability when expressed in either HUVEC *in vitro* or in zebrafish endothelial cells *in vivo* (**Figure 3**). Together, these findings suggest that *rhoaa^?^*^3^ represents a dominant-negative allele of RhoA while *rhoaa^R17^* is likely a weak dominant-negative allele.

### Rhoa and vascular integrity

Previous *in vitro* studies have implicated RHOA/ROCK function in EC barrier disruption in response to permeability inducing factors such as vascular endothelial growth factor, thrombin, histamine, and tumor necrosis factor alpha ^15, 20, 36, 80–87^. RHOA/ROCK have also been shown to mediate EC barrier enhancement in response to angiopoietin 1 and sphingosine-1-phosphate ^22–24, 88, 89^. Our zebrafish studies suggest that the *in vivo* developing cranial vasculature is exquisitely sensitive to either increased or decreased Rhoa dosage or activity (**Figure 6**). Cranial hemorrhage occurs in heterozygotes of both the stronger dominant-negative *rhoaa^?3^* mutant and the weaker *rhoaa^R17^* mutant (**Figure 1**), as well as in transgenic embryos with EC-specific expression of *rhoaa^?3^* or *rhoaa^R17^* (**Figure 4**). Embryos treated with the ROCK1/2 inhibitor Rockout (**Figure 5**) also hemorrhage, indicating that decreased RHOA/ROCK function leads to disruption of vascular integrity *in vivo*. Transgene-driven expression of ectopic wild type *rhoaa* in ECs also leads to cranial hemorrhage in zebrafish (**Figure 3**) and causes adherens junction disruption in HUVEC (**Figure 2**), suggesting that increased RHOA/ROCK function can also disrupt vascular integrity. Our combined results therefore suggest that a proper level of Rhoa/Rock signaling is required to maintain cranial vascular integrity *in vivo* under normal physiologic conditions, with either too much or too little resulting in disruption of barrier function (**Figure 6**).

**Figure 6.**
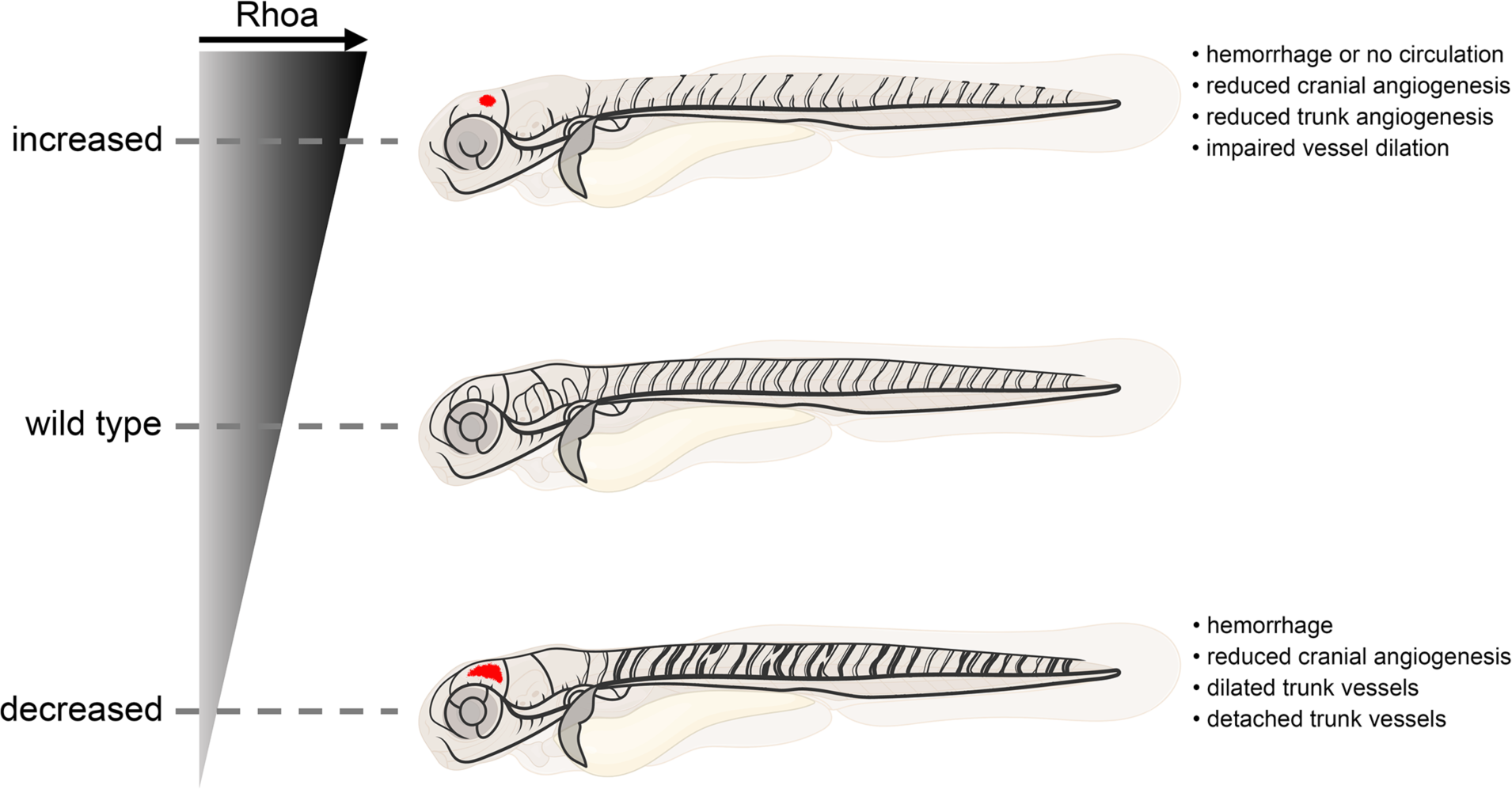
Summary of embryonic vascular defects generated by increased or decreased Rhoa signaling. Schematic diagrams illustrating the range of different vascular defects found in 52 hpf zebrafish embryos with either increased, wild type, or decreased endothelial cell levels of Rhoa activity. Embryos drawn in lateral view, anterior to the left, with blood vessels in black and cranial hemorrhage in red.

It remains unclear why the barrier-disrupting effects of increased or decreased RHOA/ROCK activity are restricted to the cranial vessels of zebrafish. In *rhoaa* mutants, EC-specific *rhoaa* transgene-expressing embryos, and Rock-inhibitor treated embryos, hemorrhage is observed in cranial, but not trunk vasculature beginning at 48 hpf. The trunk vasculature, does, however, exhibit other phenotypes such as impaired angiogenesis, vessel detachment, or vessel dilation (eg, **Figure 4R-T**). It is not clear what intrinsic molecular or physiological properties make the cranial EC barrier more sensitive to increased or decreased Rhoa activity and/or more prone to rupture than the trunk EC barrier at this developmental time point. Zebrafish embryos undergo a significant increase in peak blood pressure between 40 hpf and 60 hpf ^90^, a time when cranial vessels are still undergoing significant growth and remodeling, which could impact junctional adhesion. Trunk vessel angiogenesis (ISV growth to the DLAV) is largely complete by 40 hpf ^75^, so the differential sensitivity of cranial versus trunk vessels at this time could reflect differences in the timing of their growth and development. Alternatively, there may be intrinsic differences in the molecular and genetic profiles of cranial versus trunk ECs contributing to their differential sensitivity to gain or loss of Rhoa/Rock activity.

Interestingly, *rhoaa* mutants (*rhoaa^R17/+^*, rhoaa*^R17/^*^R17^, and *rhoaa^?3/+^* embryos) typically exhibit recovery of the EC barrier and clearance of pooled erythrocytes from the brain by 3 dpf (data not shown). This is approximately when EC barrier-protective pericytes are recruited to cranial blood vessels and the EC barrier begins to adopt a less permeable blood brain barrier (BBB) identity, characterized by increased numbers of junction proteins and reduced non-specific transcytosis events ^91–95^. Developing cranial vessels may therefore be particularly sensitive to Rhoa levels only prior to pericyte recruitment and/or BBB development.

### Rhoa and developmental angiogenesis

Previous *in vitro* studies have shown that RHOA/ROCK is required for EC migration, proliferation, and angiogenic tip cell protrusion in response to exogenous stimulation by S1P, VEGF, and/or flow ^19, 40, 80^. We find that strong suppression of Rhoa function in transgenic zebrafish with EC-specific expression of dominant-negative *rhoaa^N19^* or *rhoaa^?3^* or animals treated with ROCK1/2 inhibitor results in dramatically reduced cranial vessel angiogenesis, but has a limited effect on trunk vessel angiogenesis. The presumably weaker suppression of Rhoa function in *rhoaa^R17^* or *rhoaa^?3^* mutants, while sufficient to disrupt cranial vascular integrity (as is evident from the cranial hemorrhage in these embryos), is insufficient to disrupt cranial vessel angiogenesis. Interestingly, reduced cranial vessel angiogenesis is also noted in transgenic embryos expressing ectopic wild type *rhoaa* or constitutively-active *rhoaa^V14^* in ECs (**Figure 3**), although *rhoaa^V14^* expression might be causing EC death independently from its inhibitory effects on angiogenesis (note again our inability to maintain stable lines of *rhoaa^V14^-* expressing HUVEC and the punctate, sometimes fragmented appearance of *rhoaa^V14^*-expressing ECs in transgenic zebrafish). Together, these results suggest that, as for vascular integrity, cranial developmental angiogenesis is disrupted by too little or too much Rhoa activity.

The reasons for the more severe angiogenic defects in cranial vs. trunk vessels are unclear, although the later growth of cranial vessels compared to trunk intersegmental vessels could be a factor. Alternatively, the differential sensitivity of cranial vessels to angiogenesis defects in response to altered RHOA/ROCK could reflect intrinsic molecular differences between ECs in these different vascular beds.

### Rhoa and vascular morphogenesis

Zebrafish with altered vascular endothelial Rhoa function also exhibit defects in vascular morphogenesis and blood vessel dilation. A previous report showed that HUVEC with excess RHOA activity have impaired vacuole formation and fail to form “lumens” in *in vitro* cell culture tubulogenesis assays ^14^. EC-specific *Rhoa* knockout mice exhibit increased blood vessel lumen diameter, and EC-specific *Rhoa* knockout can partially rescue lumen formation in Ras interacting protein 1 (*Rasip1*) knockout mouse embryos, which normally exhibit increased RHOA/ROCK activity ^39^, supporting the idea that RHOA functions to inhibit or limit vessel lumenogenesis *in vivo*.

Our results are generally in line with this previously described presumptive activity for RHOA. In transgenic zebrafish with EC-specific expression of ectopic wild type *rhoaa*, vessel segments strongly expressing the transgene (as indicated by robust Tomato reporter fluorescence) are frequently narrow and undilated. In contrast, vessel segments in transgenic zebrafish with strong EC-specific expression of dominant-negative *rhoaa^N19^*, *rhoaa^?3^*, or *rhoaa^R17^* isoforms often appear excessively dilated compared to vessel segments in the same animals with less, or no transgene expression. The range of vessel dilation phenotypes observed in these individual “mosaics” strongly suggests that they are not due to reduced cardiac function and/or impaired circulation. Interestingly, zebrafish embryonic trunk vessels expressing the dominant-negative isoforms of *rhoaa* also undergo sporadic detachment (**Figure 4**). This phenotype is not accompanied by hemorrhage, suggesting that it is not simply the result of generalized EC-barrier breakdown. Future studies will be needed to define the molecular basis for this phenotype.

### Rhoa function in the vasculature

Together, our findings highlight roles for Rhoa function in vascular integrity, angiogenesis, and vascular morphogenesis during development (**Figure 6**). Too much or too little Rhoa function impairs cranial vascular integrity and cranial developmental angiogenesis. Cranial vascular integrity appears to be more sensitive to altered Rhoa gene dosage or activity than cranial angiogenesis, which apparently tolerates more substantial deviations from normal Rhoa levels or activity. Our zebrafish results largely support previous reports that RHOA functions to inhibit or limit vessel lumenogenesis. Suppressing Rhoa function in zebrafish endothelium leads to vessel dilation, while excess EC Rhoa results in undilated vessels. In addition to providing important new information on the *in vivo* roles of Rhoa in the vascular endothelium, our study highlights the power of the zebrafish as a model for genetic and experimental dissection of vascular biology in an intact animal.

## Non-standard Abbreviations and Acronyms

EC: endothelial cell
RHOA: ras homolog gene family, member A
ROCK: Rho-associated protein kinase
CCM: cerebral cavernous malformation
hpf: hours post fertilization
ENU: N-ethyl-N-nitrosourea
SSLP: Simple sequence length polymorphism
LDA: lateral dorsal aortae
PHBC: primordial hindbrain channel
CtA: cranial central artery
DA: dorsal aorta
CV: cardinal vein
ISV: intersegmental vessel
DN: dominant negative
CA: constitutively active
BBB: blood brain barrier
HUVEC: human umbilical vein endothelial cells
EGFP: enhanced green fluorescent protein
UAS: upstream activating sequence
2A: P2A viral cleavage peptide
Tg: transgenic

## Acknowledgements

The authors would like to thank members of the Weinstein laboratory for their critical comments on this manuscript. Schematics of larval zebrafish were created with BioRender.com.

## Supplemental Materials

Figures S1-S7

Table S1

Movies S1-S4

