## Supplemental Materials for "*In vivo* dissection of Rhoa function in vascular development using zebrafish"

### SUPPLEMENTAL FIGURES AND FIGURE LEGENDS

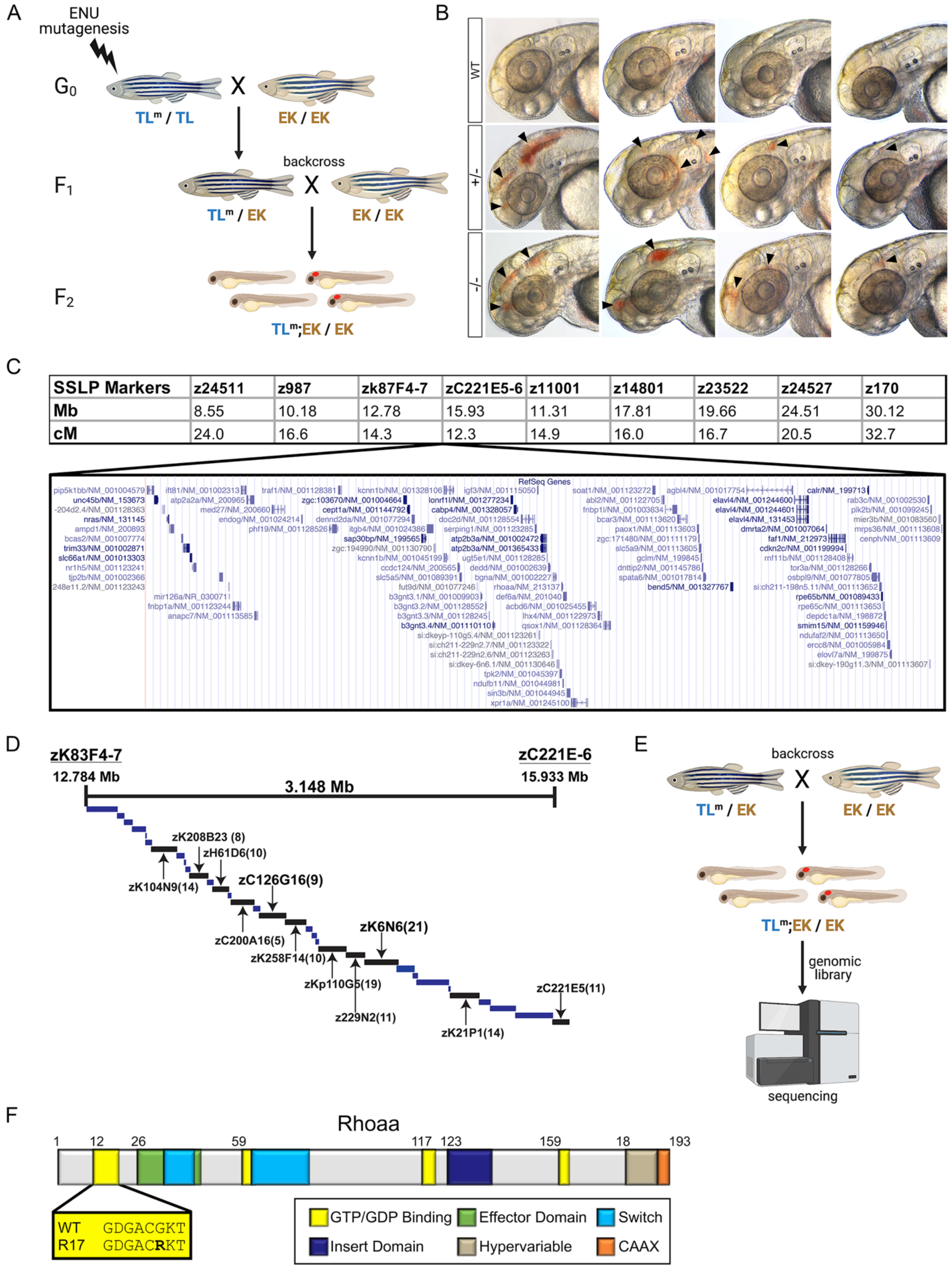

**Figure S1.** ***rhoaa*** **is mutated in** ***y172*** **mutants.** (**A**) Schematic diagram of the forward-genetic ENU mutagenesis screen used to identify dominant zebrafish mutants with intracranial hemorrhage. Tübingen Long Fin (TL) strain males were mutagenized with ENU and crossed to *Tg(fli:EGFP)*$^{y1}$ Ekkwill (EK) strain females. F1 embryos were backcrossed to EK. F2 embryos were screened for hemorrhage phenotypes. Superscript m indicates causative mutation. (**B**) Stereoscope transmitted light images of 52 hpf *y172*$^{+/-}$ incross progeny heads shown in lateral view, anterior to left. Arrowheads indicate hemorrhage. (**C**) Top: Fine mapping analyses linking the *y172* mutation to a 3.148 mega base pair (Mb) region on Chromosome 8, in between the determined SSLP markers ZK83F4-7 and ZC221E5-6. Cm denotes centimorgans. Bottom: NCBI Reference Sequence (RefSeq) database genes within the 3.148 Mb interval (Zv9/danRer7 Assembly). (**D**) Genomic clones and sequences spanning the 3.148 Mb interval, with BAC names listed. (**E**) Schematic diagram of the whole-genome sequencing approach used to identify the *y172* causative mutation. Genomic DNA libraries were constructed from 18 hemorrhaging offspring and from 18 phenotypically wild type siblings derived from a single F1 backcross pair, and both libraries were sequenced. Mutations within the critical mapped interval were filtered to include only unannotated nonsynonymous mutations. (**F**) Schematic of the 193 amino acid long Rhoaa protein, which features the same domains and motifs found in mammalian RHOA. Rhoaa contains domains involved in GTP/GDP binding (yellow), an effector domain (green), switch domains (light blue) that regulate GDP- and GTP-bound conformational changes, an insert domain (dark blue) that regulates effector binding, transforming ability, and protein stability, a hypervariable region (brown) that distinguishes RHO family members, and a CAAX domain (orange) that is required for membrane binding. Fine mapping and sequencing of *y172* mutants revealed a G-to-A substitution in Rhoaa that converts amino acid 17 in the GTP/GDP binding domain of Rhoaa from a Glycine to an Arginine (mutation bolded).

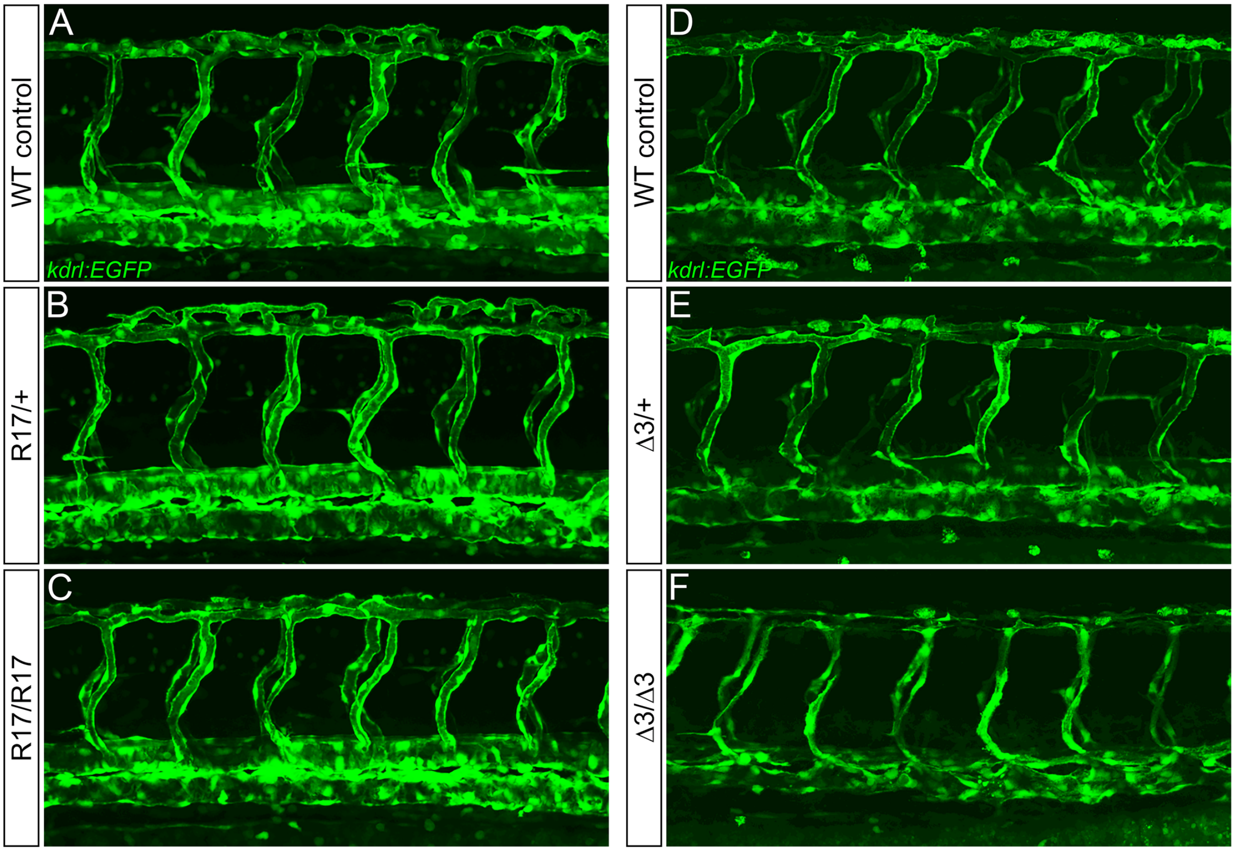


**Figure S2. Zebrafish *rhoaa* mutants exhibit normal trunk vessel patterning.** (**A-C**) Confocal images of EGFP-positive trunk vasculature in 52 hours post fertilization (hpf) *Tg(kdrl:EGFP) rhoaa*$^{R17/+}$ incross progeny shown in lateral view, anterior to left. (**D-F**) Confocal images of EGFP-positive trunk vasculature in 52 hpf *Tg(kdrl:EGFP) rhoaa*$^{\Delta 3/+}$ incross progeny shown in lateral view, anterior to left.

A

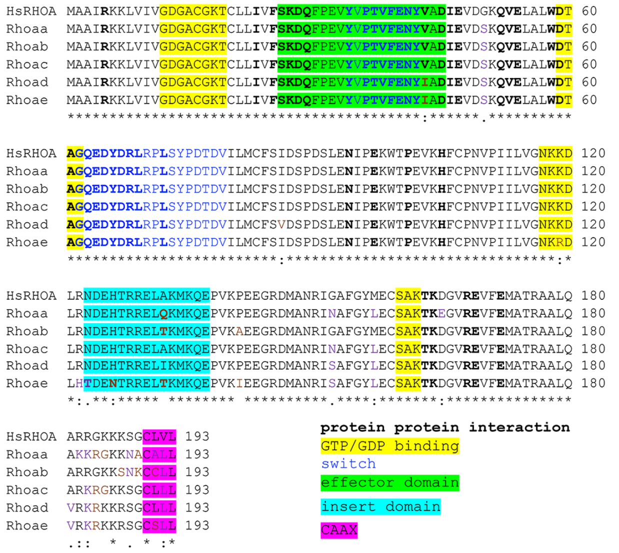


B

| Orthologue | Identity (%) |
|---|---|
| Rhoac | 96.4 |
| Rhoab | 95.9 |
| Rhoad | 94.3 |
| Rhoaa | 93.3 |
| Rhoae | 92.2 |

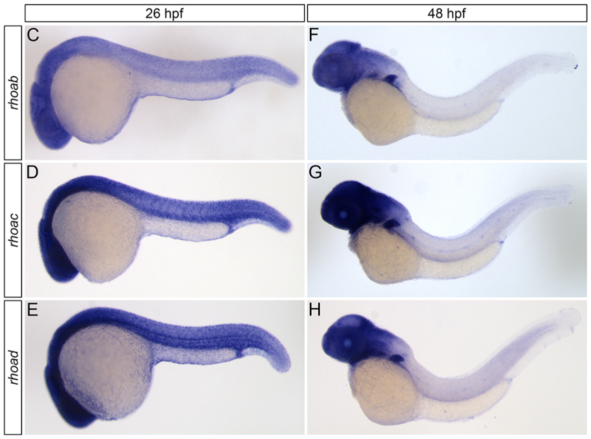


**Figure S3. Zebrafish possess several highly conserved RHOA orthologues with overlapping expression patterns.** (**A**) Amino acid sequence alignment of human RHOA (HsRHOA) and zebrafish RHOA orthologues. (**B**) Table indicating percentage of identical amino acids shared between human RHOA and zebrafish RHOA orthologues. (**C-E**) Whole-mount *in situ* hybridization analyses of zebrafish RHOA-orthologue gene expression in 26 hours post fertilization (hpf) wild type (WT) zebrafish embryos shown in lateral view, anterior to left. (**F-H**) Whole-mount *in situ* hybridization analyses of zebrafish RHOA-orthologue gene expression in 48 hpf WT zebrafish embryos shown in lateral view, anterior to left.

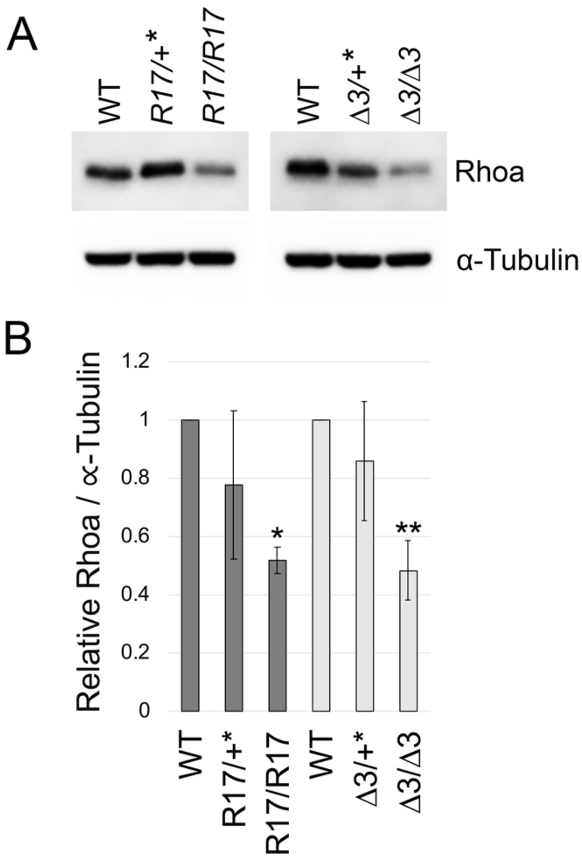


**Figure S4. Zebrafish *rhoaa* mutants possess reduced total Rhoa protein.** (**A**) Representative western blots of whole embryo protein lysates from 50 hpf zebrafish. (Left) Lysates derived from wild type incross progeny (WT), *rhoaa*$^{R17/+}$ incross progeny (*R17/+**) or *rhoaa*$^{R17/R17}$ incross progeny (*R17/R17*). *R17/+** lysate was obtained from a combination of wild type, *rhoaa*$^{R17/+}$, and *rhoaa*$^{R17/R17}$ embryos. (Right) Lysates derived from wild type incross progeny, or *rhoaa*$^{\Delta 3/+}$ incross progeny (*Δ3/+** or *Δ3/Δ3)*. *Δ3/+** lysate was obtained from a combination of wild type and *rhoaa*$^{\Delta 3/+}$ embryos. Western blots were probed with anti-RHOA antibody (top) and anti-alpha tubulin antibody (loading control; bottom). (**B**) Quantification of total Rhoa protein levels normalized to alpha tubulin loading control. Protein levels are expressed as fractions of control wild type embryo amounts. *$P$ = 0.0212 compared to WT, and **$P$ = 0.0075 compared to WT by one-way ANOVA with Tukey post hoc test. Mean ± SD is shown for each graph.

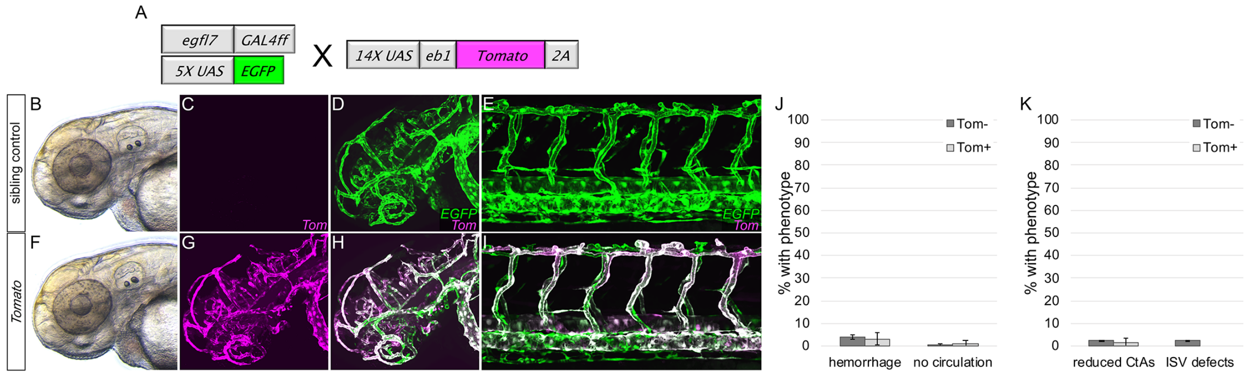


**Figure S5. Endothelial cell-specific Tomato fluorophore expression does not induce vascular integrity or patterning defects *in vivo*.** (**A**) Schematic of transgenics used for conditional Tomato fluorophore expression in zebrafish endothelial cells. (**B, F**) Stereoscope transmitted light images of 52 hpf progeny derived from a cross between *Tg(egfl7:GAL4), Tg(UAS:EGFP)* and *Tg(UAS:Tomato-2A)* fish. (**C,D,G,H**) Tomato (C,G) and Tomato/EGFP (D,H) confocal images of cranial endothelial cells in the same embryos as in panels B,F. (**E,I**) Tomato/EGFP confocal images of trunk endothelial cells in 52 hpf progeny derived from a cross between *Tg(egfl7:GAL4), Tg(UAS:EGFP)* and *Tg(UAS:Tomato-2A)* fish. (**J,K**) Quantitation of the percentage of 52 hpf Tomato-positive embryos (light grey columns) and Tomato-negative siblings (dark grey columns) with cranial hemorrhage or no circulation (J) or with reduced cranial CtAs or trunk ISV defects (K). Mean ± SD is shown for each sample.

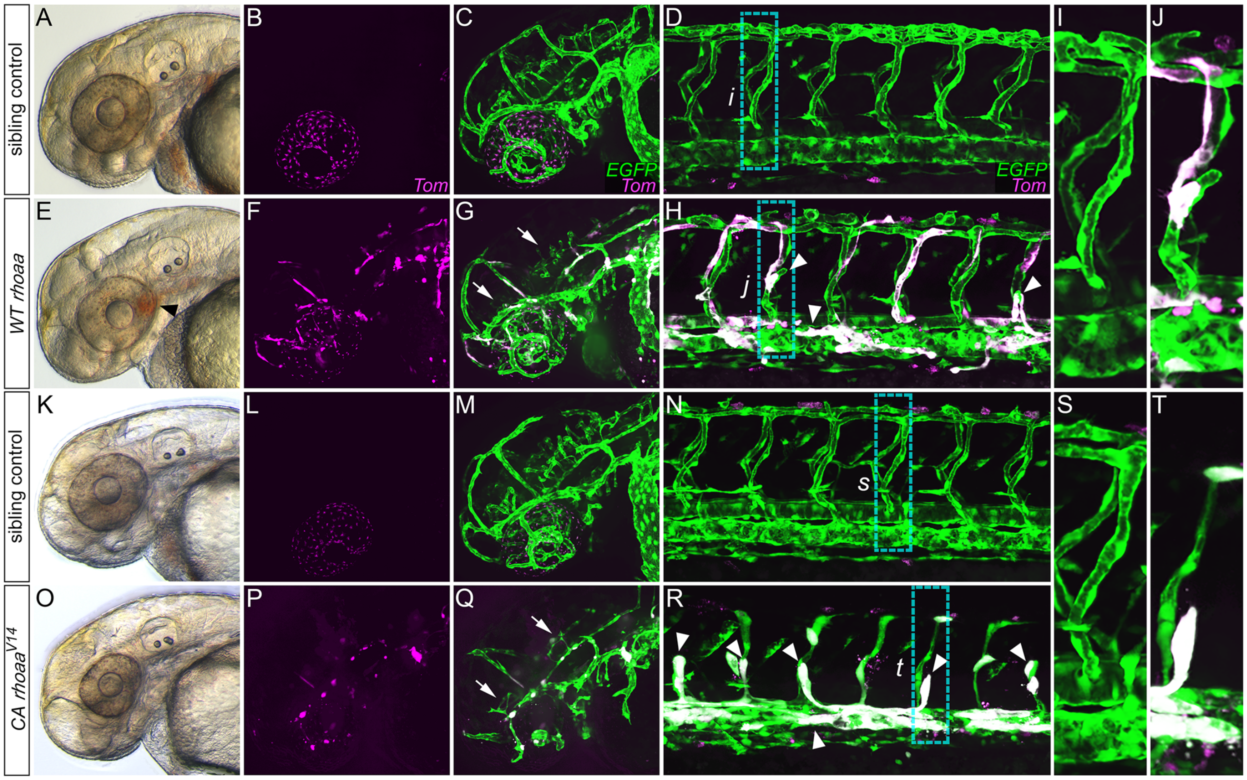


**Figure S6. Increased endothelial cell Rhoa activity induces vascular integrity and patterning defects *in vivo*.** (**A,E,K,O**) Stereoscope transmitted light images of 52 hpf progeny derived from crosses between *Tg(egfl7:GAL4), Tg(UAS:EGFP)* and *Tg(UAS:Tomato-2A-rhoaa)* (A,E) or *Tg(UAS:Tomato-2A-rhoaa$^{V14}$)* (K,O) fish. Panels A and K show non-Tomato/*rhoaa* transgene-expressing control siblings of the Tomato/*rhoaa* transgene-expressing animals in panels E and O, respectively. Black arrowhead in panel E indicates hemorrhage. (**B,C,F,G,L,M,P,Q**) Tomato (B,F,L,P) and Tomato/EGFP (C,G,M,Q) confocal images of cranial endothelial cells in the same embryos as in panels A,E,K,O. Arrows indicate reduced CtA sprouting. (**D,H,N,R**) Tomato/EGFP confocal images of trunk endothelial cells in 52 hpf progeny derived from crosses between *Tg(egfl7:GAL4), Tg(UAS:EGFP)* and *Tg(UAS:Tomato-2A-rhoaa)* (D,H) or *Tg(UAS:Tomato-2A-rhoaa$^{V14}$)* (N,R) fish. White arrowheads in panels H and R indicate ISV sprouting defects or impaired vessel dilation. (**I,J,S,T**) Magnified Tomato/EGFP confocal images of trunk vasculature in the boxed areas in panels D,H,N,R, respectively. This figure includes the same image data as in Figure 3 but with the addition of images of matched non-transgene-expressing control siblings for all crosses (i.e., new panels K-N,S).

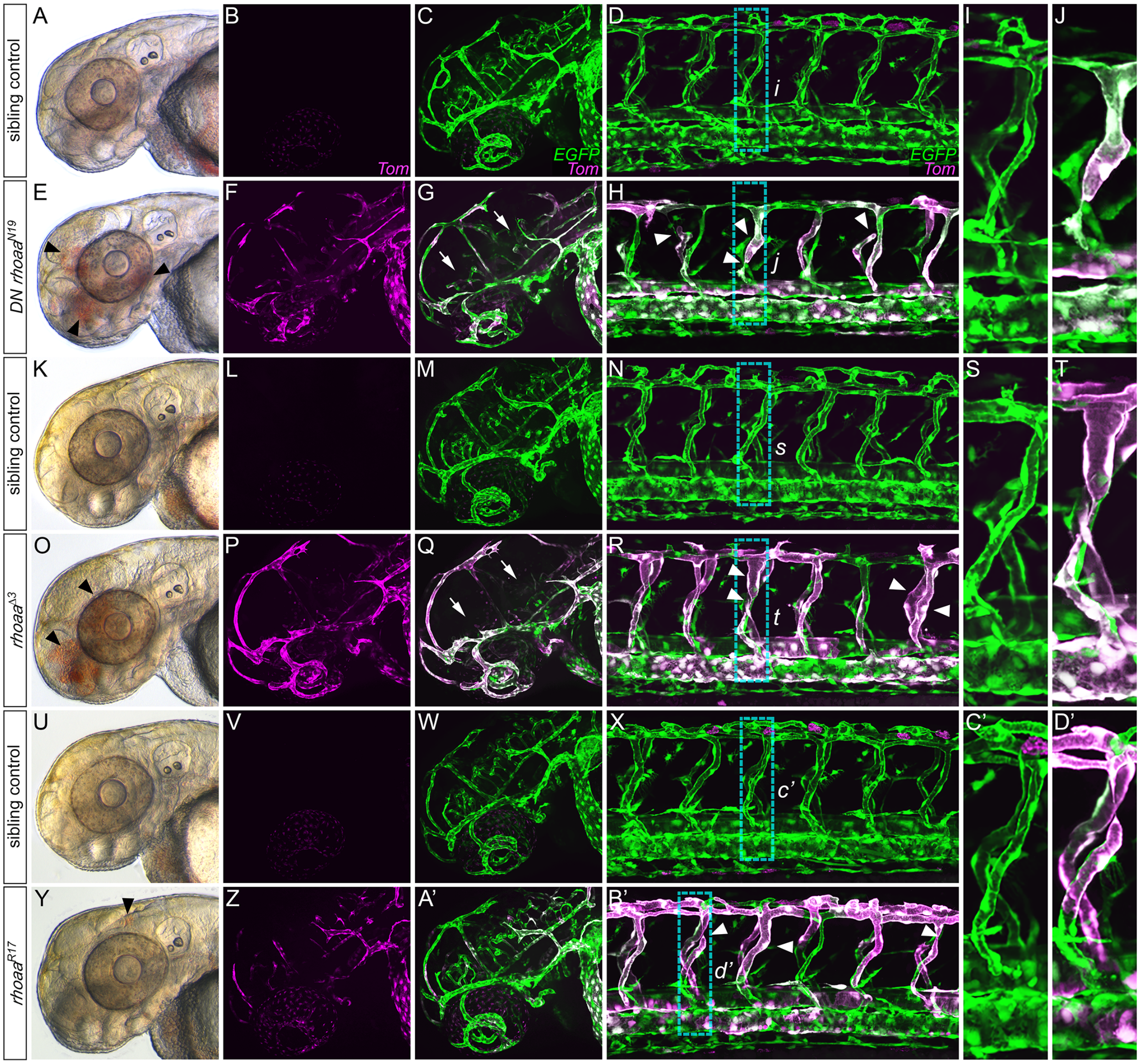


**Figure S7. Endothelial cell specific expression of dominant negative *rhoaa* induces cranial vessel integrity and patterning defects as well as trunk vessel dilation *in vivo*.** (**A,E,K,O,U,Y**) Stereoscope transmitted light images of 52 hpf progeny derived from crosses between *Tg(egfl7:GAL4), Tg(UAS:EGFP)* and *Tg(UAS:Tomato-2A-rhoaa$^{N19}$)* (A,E)*, Tg(UAS:Tomato-2A-rhoaa$^{\Delta 3}$)* (K,O), or *Tg(UAS:Tomato-2A-rhoaa$^{R17}$*) (U,Y) fish. Panels A, K, and U show non-Tomato/*rhoaa* transgene-expressing control siblings of the Tomato/*rhoaa* transgene-expressing animals shown in panels E, O, and Y, respectively. Black arrowheads in panels E,O,Y indicate hemorrhage. (**B,C,F,G,L,M,P,Q,V,W,Z,A'**) Tomato (B,F,L,P,V,Z) and Tomato/EGFP (C,G,M,Q,W,A') confocal images of cranial endothelial cells in the same embryos as in panels A,E,K,O,U,Y. Arrows in panels G and Q indicate reduced CtA sprouting. (**D,H,N,R,X,B'**) Tomato/EGFP confocal images of trunk endothelial cells in 52 hpf progeny derived from crosses between *Tg(egfl7:GAL4), Tg(UAS:EGFP)* and *Tg(UAS:Tomato-2A-rhoaa*$^{N19}$*)* (D,H)*, Tg(UAS:Tomato-2A-rhoaa*$^{\Delta 3}$*)* (N,R), or *Tg(UAS:Tomato-2A-rhoaa*$^{R17}$) (X,B') fish. Arrowheads in panels H,R,B' indicate ISV dilation, detachment, or growth defects. (**I,J,S,T,C',D'**) Magnified Tomato/EGFP confocal images of trunk vasculature in the boxed areas in panels D,H,N,R,X,B', respectively. This figure includes the same image data as in Figure 4 but with the addition of images of matched non-transgene-expressing control siblings for all crosses (i.e., new panels K-N,S,U-X,C').

### SUPPLEMENTAL TABLES

### Table S1

**To make riboprobes for *in situ* hybridization**

| Purpose | Primer Name | Sequence |
|---|---|---|
| Amplify rhoaa riboprobe sequence from cDNA | rhoaa-pF | TAGTTACTACCCTACATCGCTTATGC |
| | rhoaa-pR | GGAAATCCAAAGATGCACTGGTATAC |
| Amplify rhoab riboprobe sequence from cDNA | rhoab-pF | GAACTGACCAAGATGAAGCAGGAG |
| | rhoab-pR | GTTTACAAATACGGTTTCCAGTCCTG |
| Ampify rhoac riboprobe sequence from cDNA | rhoac-pF | AAAATGAAACAGGAGCCGGTAAAG |
| | rhoac-pR | CAGTATCGCTAAATGGTTTAGATCCC |
| Amplify rhoad/rhocb riboprobe sequence from cDNA | rhoad-pF | TGTTCAGTAAAGATCAGTTCCCTGAG |
| | rhoad-pR | CTGTAAACCAGAATCGAAAGAGACAG |
| Amplify rock1 riboprobe sequence from cDNA | rock1-pF | AAATCCTCTTTTACAATGACGAGCAG |
| | rock1-pR | AATGTGGGAAAATAGAGAAAGATGGC |
| Amplify rock2a riboprobe sequence from cDNA | rock2a-pF | CAGTTAACCTTGTATTGGGATTCTGG |
| | rock2a-pR | GCAAATGTTATACAGGGTCTCCAAAC |

**To make p14XUAS-Tom-2A-rhoaa constructs**

| Purpose | Primer Name | Sequence |
|---|---|---|
| Amplify rhoaa coding sequence | UAS-rhoaa-F | AGACGTGGAGGAGAACCCTGGACCT**ATGGCTGCGATTCGTAAGAAGCTTG** |
| | UAS-rhoaa-R | AGTTAACGGTGGCTGAGACTTAATT**CTATAGCAAAGCGCAGGCATTCTTC** |
| Amplify UAS vector | UAS-TomVec-F | **AATTAAGTCTCAGCCACCGTTAACTG** |
| | UAS-TomVec-R | **AGGTCCAGGGTTCTCCTCCAC** |
| G14V rhoaa site-directed mutagenesis | rhoaa-G14V-F | GTTGGAGATG**T**AGCCTGTGGAAAG |
| | rhoaa-G14V-R | GATTACAAGCTTCTTACGAATC |
| T19N rhoaa site-directed mutagenesis | rhoaa-T19N-F | TGTGGAAAGA**AC**TGTTTACTCATCGTTTTCAGTAAAG |
| | rhoaa-T19N-R | GGCTCCATCTCCAACGAT |

**To genotype rhoaa∆3 CRISPR zebrafish mutants**

| Purpose | Primer Name | Sequence |
|---|---|---|
| Genotyping PCR primers | rhoaaM13Ftail-F | TGTAAAACGACGGCCAGT**TTCCACCTGCCTCCTGTAATTATTG** |
| | rhoaaPigtail-R | GTGTCTT**CAGCGACATAGTTCTCAAACACTG** |
| | FAM-M13 primer | /56-FAM/TGTAAAACGACGGCCAGT |

**To make pCDH-EF1-Tom-2A-Rhoaa-(sCMV-GFP-T2A-PURO) constructs**

| Purpose | Primer Name | Sequence |
|---|---|---|
| Amplify pCDH without PGK | pCDH-minusPGK-F | AGAGGCCACTTGTGTAGCGCCAAG |
| | pCDH-minusPGK-R | CTGCAGCCCAAGCTTACCATGGAG |
| Amplify sCMV from pCS2+ | sCMV-F | GCTCTCGTCGCTCTCCATGGTAAGCTTGGGCTGCAG**GCTCCGACGTCCCCAGGCAGAATG** |
| | sCMV-R | CTTGGCGCTACACAAGTGGCCTCTGGCCTCGCACAC**GACCATAGCCAATTCAATATGGCG** |
| Amplify rhoaa for insertion into pCDH | pCDH-rhoaa-F | GCTGCGATTCGTAAGAAGCTTGTAATCG |
| | rhoaaBamHI-R | GTAATCCAGAGGTTGATTGTCGACGCGGATCC**CTATAGCAAAGCGCAGGCATTCTTC** |
| Amplify pCDH for rhoaa or Tomato insertion | pCDH-rhoaa-F | GGATCCGCGTCGACAATCAACCTCTGGATTAC |
| | pCDH-R | GAATTCGCTAGCTCTAGATCACGACACC |
| Amplify pCDH-rhoaa for Tomato insertion | pCDH-rhoaa-F | GCTGCGATTCGTAAGAAGCTTGTAATCG |
| | pCDH-Tom-R | CGAATTCGCTAGCTCTAGATCACGACACC |
| Amplify Tomato for insertion into pCDH (with rhoaa) | pCDH-Tom-F | GTCGTGATCTAGAGCTAGCGAATTCGCCACC**ATGGTGAGCAAGGGCGAGGAGGTC** |
| | Tom-rhoaa-R | TTACAAGCTTCTTACGAATCGCAGCAGCGCT**CTTGTACAGCTCGTCCATGCCGTACAGG** |
| Amplify Tomato for insertion into pCDH (without rhoaa) | pCDH-Tom-F | GTCGTGATCTAGAGCTAGCGAATTCGCCACC**ATGGTGAGCAAGGGCGAGGAGGTC** |
| | Tom-pCDH-R | GTAATCCAGAGGTTGATTGTCGACGCGGATCC**CTTGTACAGCTCGTCCATGCCGTACAGG** |

#### LEGENDS FOR VIDEO FILES

**Movie S1. Live imaging of heartbeat and blood flow in wild type and *rhoaa*$^{\Delta3/\Delta3}$ embryos.** Videos of 52 hpf wild type and *rhoaa*$^{\Delta3/\Delta3}$ embryos shown in lateral view, anterior to left. Blood flow through the beating heart, cranial and trunk vasculature is readily observed in the wild type but not *rhoaa*$^{\Delta3/\Delta3}$ embryo.

**Movie S2. Live imaging of heartbeat and blood flow in wild type embryos and embryos with EC-specific constitutively active *rhoaa*$^{V14}$ expression.** Videos of cranial and trunk circulation in 52 hpf progeny derived from a cross between *Tg(egfl7:GAL4), Tg(UAS:EGFP)* and *Tg(UAS:Tomato-2A-rhoaa*$^{V14}$*)* fish, shown in lateral view, anterior to left. Confocal images of corresponding transgene expression accompany each video. Blood flow through the beating heart, cranial and trunk vasculature is readily observed in wild type sibling control, but not the embryo with *Tg(UAS:Tomato-2A-rhoaa*$^{V14}$*)* transgene expression.

**Movie S3. Cranial and trunk blood vessel morphology in an embryo with EC-specific constitutively active *rhoaa*$^{V14}$ expression and its sibling control.** Video rotations of fluorescent 3D confocal z-stacks highlighting the cranial and trunk vessels of 52 hpf progeny derived from a cross between *Tg(egfl7:GAL4), Tg(UAS:EGFP)* and *Tg(UAS:Tomato-2A-rhoaa*$^{V14}$*)* fish. Note impaired dilation of trunk vessels and rounded morphology of cranial blood vessel ECs in embryo with *rhoaa*$^{V14}$ expression (Tomato-positive), compared to its sibling control (Tomato-negative).

**Movie S4. Trunk vasculature in an embryo with EC-specific dominant negative *rhoaa*$^{N19}$ expression.** Video rotation of a fluorescent 3D confocal z-stack highlighting trunk intersegmental vessel detachment in a 52 hpf *Tg(egfl7:GAL4), Tg(UAS:EGFP)* embryo with mosaic *Tg(UAS:Tomato-2A-rhoaa*$^{N19}$*)* transgene expression. Trunk vasculature is shown in lateral view with anterior to right.
